# Chronic alcohol exposure drives inflammaging and transposon derepression in hematopoietic stem progenitor cells

**DOI:** 10.64898/2025.12.25.696246

**Authors:** Ridzky A. A. Yuda, Habin Bea, Valerie Kellett, Jungwoo Kim, Fan Yang, Haruna B. Choijilsuren, Yejie Park, Yaojie Fu, Zewen Ha, Juhye Choi, Li Z. Luo, Zhaoli Sun, Bin Gao, Sangmoo Jeong, Linda M. S. Resar, Moonjung Jung

## Abstract

Chronic alcohol use can cause pancytopenia and diminished immune responses against pathogens. However, its underlying molecular mechanisms remain unclear. Furthermore, whether chronic alcohol consumption directly induces inflammation in human hematopoietic stem progenitor cells (HSPCs) or whether it affects aging hematopoiesis differently is unknown. To examine how chronic alcohol use influences HSPCs, we performed single-cell RNA-seq in murine and human HSPCs and single-cell ATAC-seq in aged murine HSPCs following alcohol exposure. In the native murine bone marrow, chronic alcohol exposure primed HSPCs to differentiate into myeloid cells and to exhibit heightened inflammation, DNA damage, and epigenetic reactivation of transposable elements (TEs) in an age-dependent manner. Alcohol-exposed aged long-term hematopoietic stem cells (LT-HSCs) displayed increased chromatin accessibility at TE-containing loci correlated with aberrant TE transcription. This transposon derepression was associated with the accumulation of dsRNAs in aged bone marrow cells, and activation of innate immune pathways, perpetuating HSC inflammaging. Furthermore, we identified two epigenetically distinct LT-HSC clusters, LT-HSC1 and LT-HSC2, with the LT-HSC2 cluster expanding in response to chronic alcohol consumption, resembling activated HSCs. In xenotransplanted human HSPCs, chronic alcohol feeding resulted in a significant myeloid bias, heightened inflammation, upregulation of double-stranded RNA (dsRNA) sensors, activation of type I interferon responses, and increased expression of endogenous retroviruses. Despite these molecular alterations, we did not observe a decrease in long-term repopulation capacity in either human or murine HSCs. This suggests that HSC function may recover following alcohol cessation. However, previous chronic alcohol exposures imprint murine HSPCs to exhibit long-term myeloid bias and reduced cell cycle entry upon bacterial LPS challenge. Our data illuminate potential interactions between alcohol and aging that can reinforce inflammaging and epigenetic dysregulation in HSPCs.

**Keypoints:**

1. Aging perpetuates alcohol-induced myeloid bias, inflammation, DNA damage, and TE upregulation in murine HSPCs
2. Prior chronic alcohol consumption does not affect long-term repopulation but causes persistent myeloid bias and inefficient stress responses after LPS challenge
3. Chronic alcohol consumption alters chromatin accessibility in TE-overlapping regions
4. Chronic alcohol consumption promotes myeloid bias, inflammation, and upregulation of endogenous retroviruses in xenotransplanted human HSPCs.

**Graphical Abstract:** 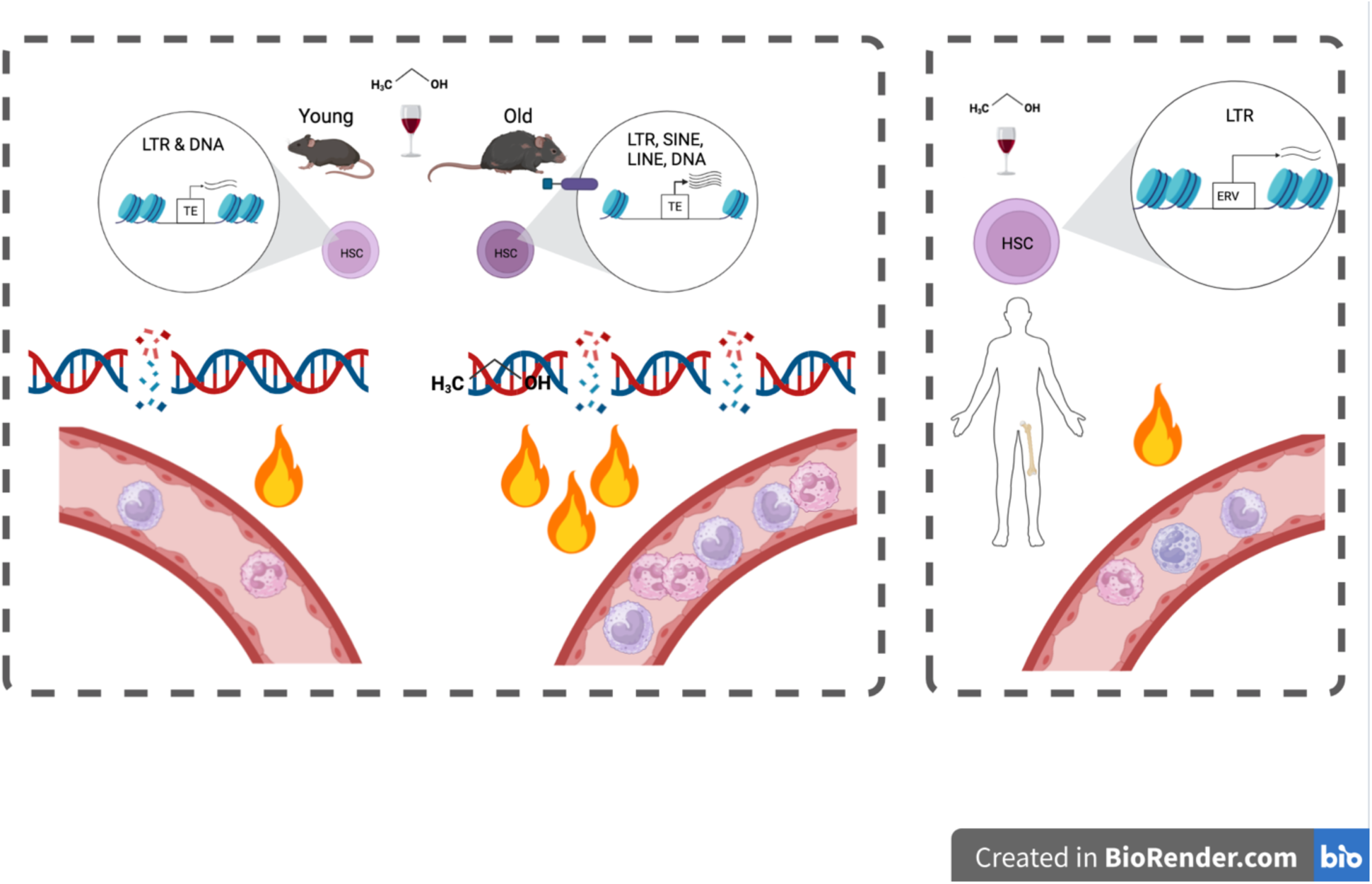

## Introduction

About 2.6 million deaths were caused by alcohol consumption worldwide in 2019^1^. Approximately 10% of the U.S. population suffers from alcohol use disorder (AUD)^2^, associated with increased morbidity and mortality from liver disease, cancer, infection, and/or bone marrow suppression^3^. Furthermore, a rapidly growing number of older adults (ages ≥ 65) with heavy or binge alcohol use is of great public health concern^4^, and it is associated with a higher alcohol-related mortality^4^. Therefore, there is a growing unmet need to better understand the molecular consequences of chronic alcohol consumption in older individuals.

Ethanol and its derivative metabolites promote DNA damage^5^ and alter cellular metabolism in the liver^6–8^, brain^9^, and hematopoietic system^10–13^. Chronic alcohol consumption causes thrombocytopenia, anemia, leukopenia, or even pancytopenia^10–12^, indicating impaired hematopoiesis. In macaques, chronic alcohol consumption increases inflammation and hematopoietic stem cells’ (HSCs) differentiation into monocytes^14^ and osteoclasts^15^, while blocking erythroid differentiation^16^. However, the molecular underpinnings of heightened inflammation and myeloid bias are yet to be fully understood. Aging primes HSCs to upregulate genes involved in cell adhesion, ribosomal biogenesis, and proliferation, and to downregulate DNA repair and epigenetic maintenance genes^17,18^. These alterations increase DNA damage^19,20^, inflammation, and myeloid bias in HSCs, key features of inflammaging^21^. However, it is unknown whether alcohol accelerates inflammaging in HSCs. Furthermore, a recent preprint reports two distinct HSC subsets during aging and inflammation^22^. Whether chronic alcohol consumption and aging differentially affect the two HSC subsets remains unclear.

Transposons or transposable elements (TEs) are mobile repetitive DNA sequences in the genome^23^. TEs are classified into long-terminal repeat (LTR), long- and short-interspersed nuclear element (LINE and SINE) retrotransposons, and DNA transposons^23^. Chromatin-modifying enzymes, including *krüppel*-associated box zinc-finger proteins (KRAB-ZFP)^24,25^, P-element induced Wimpy testis (PIWI)^26^, and DNA methyltransferases (DNMTs) repress TEs^27–29^, and their loss leads to TE reactivation during aging^18^. TE-derived nucleic acids can activate nucleic acid sensors, such as cGAS-STING or double-stranded RNA (dsRNA) sensors, prompting cells to trigger interferon responses and enter premature senescence^30^. Furthermore, TEs also carry transcription factor (TF) binding sites^31^, such as CCCTC-binding factor (CTCF), which regulates gene expression during inflammation^32^. However, whether alcohol-induced DNA damage causes epigenetic changes, leading to TE derepression and upregulation of inflammatory genes, is poorly understood.

Here, we report how chronic alcohol consumption influences murine and human HSPCs and how aging modifies its effects. In the native bone marrow niche, chronic alcohol consumption exacerbated inflammatory and aging-associated features, including TE derepression in aged murine HSPCs. Notably, long-term repopulation was maintained, but a myeloid bias and inefficient stress response were persistent in previously alcohol-exposed HSCs. Importantly, chronic alcohol exposure elicits inflammatory signaling in xenotransplanted human HSPCs, characterized by activation of dsRNA sensors, type I interferon responses, and upregulation of endogenous retroviruses. Together, these findings reveal that alcohol and aging synergistically reinforce inflammation and epigenetic dysregulation in HSPCs.

## Methods

### Mice and Husbandry

We used 5 to 6-week-old female NBSGW mice (JAX #026622), 2-month-old C57BL/6NJ (JAX #005304), 21 to 22-month-old C57BL/6 mice (National Institute of Aging Aged Rodent Colonies), 6 to 8-week-old female C57BL/6J (CD45.2; JAX #000664), and B6.SJL-*Ptprc^a^ Pepc^b^*/BoyJ (CD45.1; JAX #002014). Both sexes were used unless otherwise stated. Mice were housed in the specific-pathogen-free animal facility in the Miller Research Building.

### Study Approvals

The use of deidentified healthy volunteer BM cells was approved by Johns Hopkins Medicine Institutional Review Board. All animal experiments were approved by Johns Hopkins University Animal Care and Use Committee.

### Alcohol feeding experiments

A modified NIAAA alcohol feeding protocol^33^ was done without binge. Lieber-DeCarli isocaloric control and ethanol liquid diets were obtained from Bio-Serv. Mice were acclimated to the liquid diet with a gradual dose escalation until reaching 5% v/v ethanol. The 5% alcohol feeding was conducted for 8 weeks in NBSGW mice and for 4–5 weeks in C57BL/6 mice. Survival and consumption of the liquid diet were monitored daily, and body weight was monitored every 2-3 days. Alcohol concentration was reduced to 4% v/v for old female mice at the 4^th^ week, when premature death was observed.

### Cell harvest and flow-cytometry analysis

Cells were isolated from peripheral blood (PB) obtained via retroorbital bleeding, from crushed spleens, and from bone marrow collected from the tibiae, femurs, pelvic bones, and humeri. Alveolar macrophages were obtained from bronchoalveolar lavage^34,35^. Cells were stained with antibody panels as described in the Supplemental Methods. Data were acquired using Aurora Cytek and analyzed using FlowJo v10 (TreeStar).

### dsRNA immunofluorescence staining

Detection of dsRNAs in murine alveolar macrophages, human leukemia cell lines, and human BM CD34^+^ cells was done by immunofluorescence staining with J2 antibody (1:1000, Scicons, 10010200). Slides were imaged using EVOS M5000. Procedures are detailed in the Supplemental Methods.

### Statistical and data analysis

Results were analyzed using GraphPad Prism v10, with Student’s t-test or one-way ANOVA followed by Tukey’s multiple comparison test, as indicated in the figure legends.

## Results

### Chronic alcohol consumption perpetuates myeloid bias in the aged hematopoietic system

To understand the effects of chronic alcohol consumption on young and old unperturbed hematopoiesis, we fed young and old wild-type C57BL/6 mice with a 5% alcohol (v/v) or control liquid diet for 4-5 weeks. We had 4 groups: young control (YC), young alcohol (YA), old control (OC), and old alcohol (OA) (**Figure 1A, S1A**). We achieved a mean blood ethanol concentration (BEC) of approximately 30-40 mM (**Figure S1B**). Chronic alcohol consumption did not significantly alter the body weight between the age-matched groups (**Figure S1C**). Median survival was slightly reduced in alcohol groups due to premature deaths in females, with the strongest reduction observed in old female mice, corroborating the reported greater harm of alcohol in females^36,37^ (**Figure S1D**).

**Figure 1.**
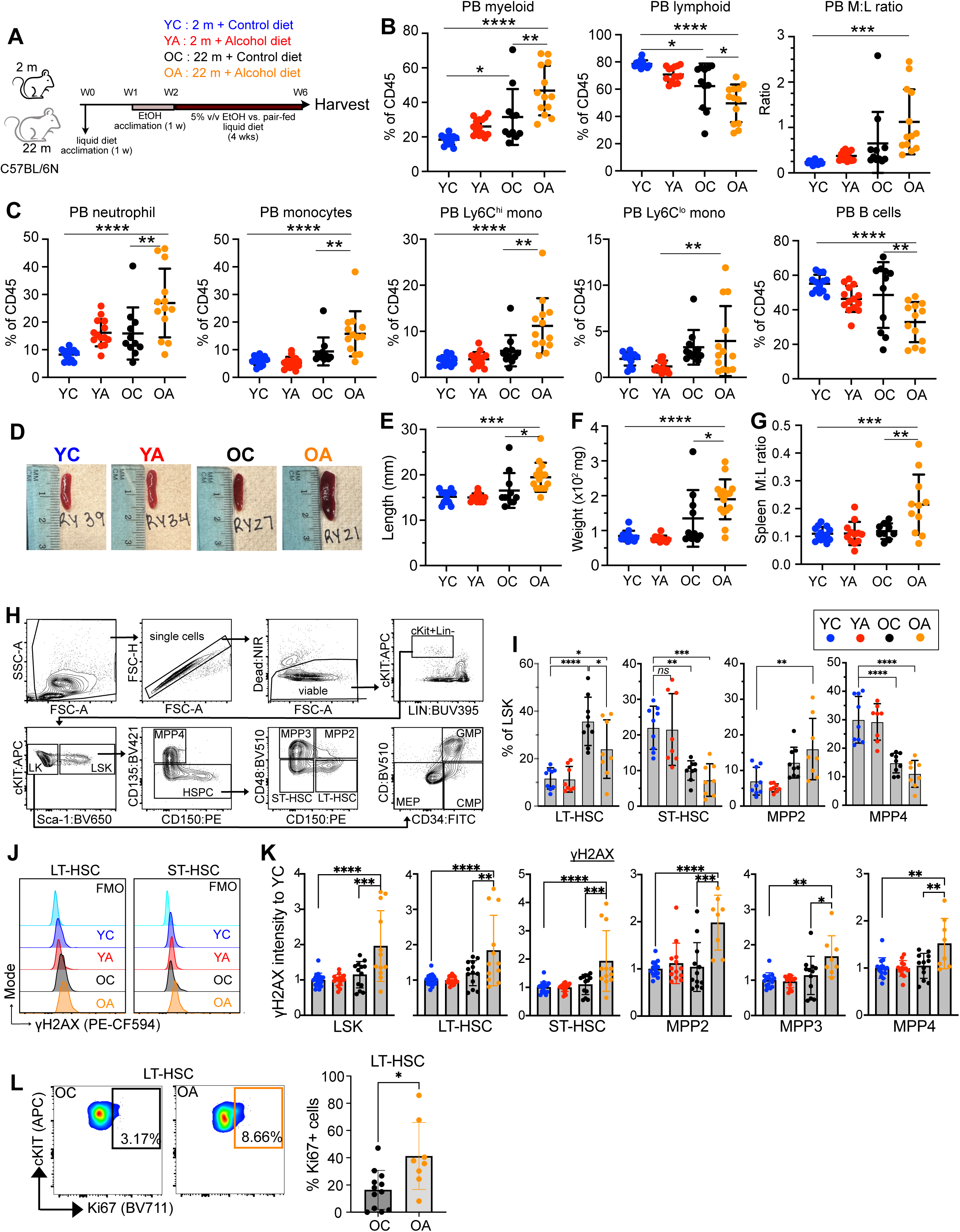
Chronic alcohol consumption caused significant myeloid bias and DNA damage in an age-dependent manner. (A) Experimental scheme of chronic alcohol feeding in young and old mice. Following a 1-week liquid diet and a 1-week alcohol diet acclimatization, the mice were given 5% v/v alcohol or a pair-fed control diet for 4 weeks. (B) Percent of myeloid and lymphoid cell subpopulations within CD45+ PB cells and PB myeloid to lymphoid (M:L) ratio, showing a significant increase between OA vs. YC, but not between OC vs. YC mice at the end of alcohol feeding (n = 11-14 mice/group; pool of 3 experiments). (C) Percent of neutrophil, total monocytes, Ly6C^hi^ and Ly6C^low^ monocytes, and B cells within PB CD45+ cells (n = 11-14 mice/group; pool of 3 experiments). (D) Representative spleen pictures from 4 different groups. (**E-F**) Quantification of spleen length (**E**) and weight (**F**) at the end of alcohol feeding (n = 11-15 mice/group; pool of 3 experiments). (G) M:L ratio in spleen (n = 10-13 mice/group; pool of 3 experiments). (H) Gating strategy to identify bone marrow HSPC subpopulations within LSK and LK compartments. (I) Proportion of LT-HSC, ST-HSC, MPP2, and MPP4 within LSK (n = 8-9 mice/group; pool of 2 experiments). (J) Representative histogram plots showing γH2AX intensity measured in LT-HSC and ST-HSC, comparing 4 groups. (K) Quantification of γH2AX intensity in the LSK, LT-HSC, ST-HSC, and MPP2 from alcohol- and control diet-fed young and old mice (n = 12-18 mice/group; pool of 4 experiments). (L) Representative flow-cytometry plots showing % Ki67+ LT-HSC in alcohol- and control diet-treated old mice (n = 8-12 mice/group; pool of 3 experiments). All data were analyzed using one-way ANOVA followed by Tukey’s multiple comparisons, except for **L**, which was analyzed using an unpaired Student t-test. *p < 0.05; **p < 0.01; ***p < 0.001; ****p < 0.0001. Bars indicate mean ± SD, dots represent individual mice. PB, peripheral blood; LSK, lineage-Sca1+ cKit+; LK, lineage-cKit+; LT-HSC, long-term HSC; ST-HSC, short-term HSC; MPP, multipotent progenitor.

We witnessed a significantly increased proportion of myeloid cells and a significantly decreased proportion of lymphoid cells in old mice, resulting in a myeloid bias, as expected with aging (**Figure 1B**). Myeloid bias became more striking with alcohol consumption, suggesting an exacerbated response to alcohol in aged mice. Neutrophil and monocyte populations increased significantly in the OA but not in the YA group, as compared with age-matched controls (see **Figure S1E** for gating strategy). Furthermore, OA mice showed elevated Ly6C^high^ inflammatory monocytes, along with reduced lymphoid cells (**Figure 1C, S1F**). Importantly, the alcohol exposure caused splenomegaly (**Figure 1D-F**) and induced myeloid bias in the spleen of OA mice (**Figure 1G, S1G**). Chronic alcohol consumption reduced hemoglobin, hematocrit, RBC, and platelet counts in OA but not in YA mice compared to YC mice (**Figure S1H**). Taken together, chronic alcohol consumption and aging synergistically heightened myeloid bias in the aging hematopoietic system.

### Chronic alcohol consumption induces DNA damage and cell-cycle entry in aged HSPCs

Next, we investigated whether aged HSPCs are more vulnerable to chronic alcohol exposure. A gating strategy from the Passegué lab was used to identify HSPC populations in Lin^-^ Sca-1^+^ cKit^+^ (LSK) and Lin^-^ Sca-1^-^ cKit^+^ (LK) compartments (**Figure 1H**)^38^. Chronic alcohol consumption did not change the proportion of LT-HSC, ST-HSC, or MPP4 in young mice (**Figure 1I**). However, alcohol exposure resulted in a relative contraction of the LT-HSC that was expanded by aging, and alcohol exposure and aging synergistically expanded MPP2 and MEP populations (**Figure 1I, S2A**). Consistent with the reduced alcohol metabolism^4,39^ and DNA repair capacity associated with aging^17,40^, chronic alcohol exposure significantly increased γH2AX intensity only in old but not in young LT-HSCs, ST-HSCs, MPP2/3/4, and LK subpopulations (**Figure 1J-K, S2B**). Furthermore, OA mice demonstrated a significantly higher proportion of cycling (Ki67+) LT-HSCs, ST-HSCs, and MPP2/3/4 (**Figure 1L, S2C**), indicating increased cell-cycle entry due to replicative stress from alcohol exposure and aging.

### Chronic alcohol consumption induces persistent myeloid bias and inefficient stress responses in HSPCs

We next asked whether prior chronic alcohol consumption reduced HSPC long-term repopulation capacity and differentiation potential. To test this, total BM cells from mice (CD45.2) exposed to a 4-5 week alcohol or control diet were co-transplanted in a 1:1 ratio with competitor BM cells (CD45.1) into lethally irradiated recipients (CD45.2) (**Figure 2A**). Engraftment was monitored by serial flow cytometry without further alcohol treatment (see **Figure S2D** for gating strategy). Chronic alcohol consumption slightly reduced engraftment of young HSCs at 1° BMT, whereas it caused fluctuation yet comparable engraftment between the OC and OA groups (**Figure 2B-C**). At the final 2° BMT, the engraftment of control- and alcohol-exposed HSCs was comparable in both age groups (**Figure 2B-C**). Interestingly, reduced B cells and increased myeloid cells (neutrophils, Ly6C^high^ and Ly6C^low^ monocytes) were persistent in OA mice at final 1° BMT, which is 20 weeks after the end of alcohol feeding, but then faded away during the 2° BMT (**Figure 2D**). Inefficient cell-cycle entry or a blunted response to LPS are well-described characteristics of aged LT-HSC^41,42^. We also challenged mice with LPS, followed by an EdU chase at two separate time points, to examine HSPCs’ ability to respond to an inflammatory stimulus. Interestingly, prior alcohol exposure (4-6 weeks earlier (**Figure 2F**) and 30-32 weeks earlier (**Figure 2G**)) resulted in decreased LPS-induced cell cycle entry in HSPCs compared to their age-matched controls (**Figure 2E-G**). These results suggest that chronic alcohol consumption causes persistent myeloid bias and inefficient cell-cycle entry in response to stress, which lasts for a long time after alcohol cessation.

**Figure 2.**
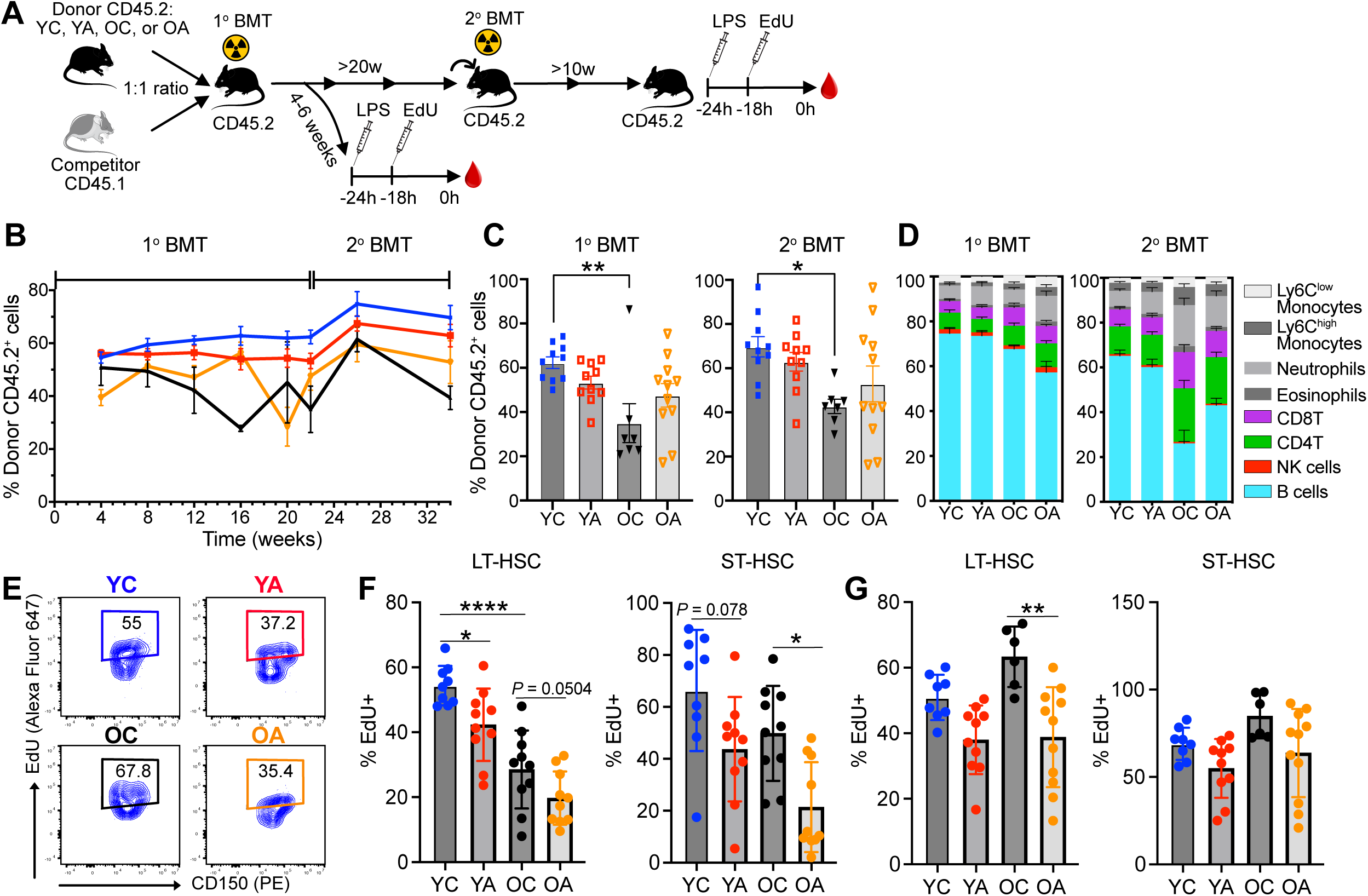
Chronic alcohol consumption alters murine hematopoietic stem cells with long-term consequences. (A) Diagram of serial bone marrow reconstitution experiments with the absence of alcohol feeding. (B) Dynamics of donor CD45.2 contribution in PB of recipient mice at primary and secondary BMT (n = 7-11 mice/group; pool of 2 experiments). (C) Donor CD45.2 contribution at 22 weeks post-1° BMT and at 10 weeks post-2° BMT measured in PB. Donor CD45.2 engraftment was significantly reduced in OC compared with YC (n = 7-11 mice/group; pool of 2 experiments). (D) Lineage contribution in PB within total CD45.2 was measured at 22 weeks post-1° BMT and at 10 weeks post-2° BMT (n = 7-11 mice/group; pool of 2 experiments) (E) Gating strategy of EdU-positive cells in LT-HSCs. (F) Proportion of EdU-positive cells in LT-HSCs and ST-HSCs after LPS challenge at 4-6 weeks post-1° BMT (n = 9-10 mice/group; pool of 2 experiments). (G) Proportion of EdU-positive cells in LT-HSCs and ST-HSCs after LPS challenge at 10-12 weeks post-2° BMT (n = 6-11 mice/group; pool of 2 experiments). All data were analyzed using one-way ANOVA followed by Tukey’s multiple comparison. All statistically significant levels were shown as * *p* < 0.05, ** *p* < 0.01, *** *p* < 0.001, **** *p* < 0.0001.

### Chronic alcohol consumption increases inflammation and alters age-associated transcriptional programs in HSPCs

To identify chronic alcohol-induced transcriptional changes in HSPCs, we performed 10X Genomics scRNA-seq from flow-sorted murine bone marrow LSK cells. Fifteen LSK clusters were identified, including the LT-HSC cluster with high *Procr*, *Pdzk1ip1,* and *Mecom* expressions and the highest stem cell module score^43^ (**Figure 3A**, **S3A-B**). The MPP2 cluster exhibited high expression of proliferative genes, and the MPP3 cluster showed a myeloid bias similar to previously described (**Figure S3B**)^5,6^. Furthermore, the MPP4 cluster was also identified based on *Flt3*, *Mn1,* and *Bcl2l11* expression. Chronic alcohol consumption reduced LT-, ST-HSC, and MPP4 clusters, whereas it expanded MPP2, MPP3, MDP, GMP, and MEP clusters in OA mice (**Figure 3B**). Unexpectedly, we identified an unknown cluster that was significantly enriched in the OA mice and lacked specific lineage-defining markers (**Figure S3B**).

**Figure 3.**
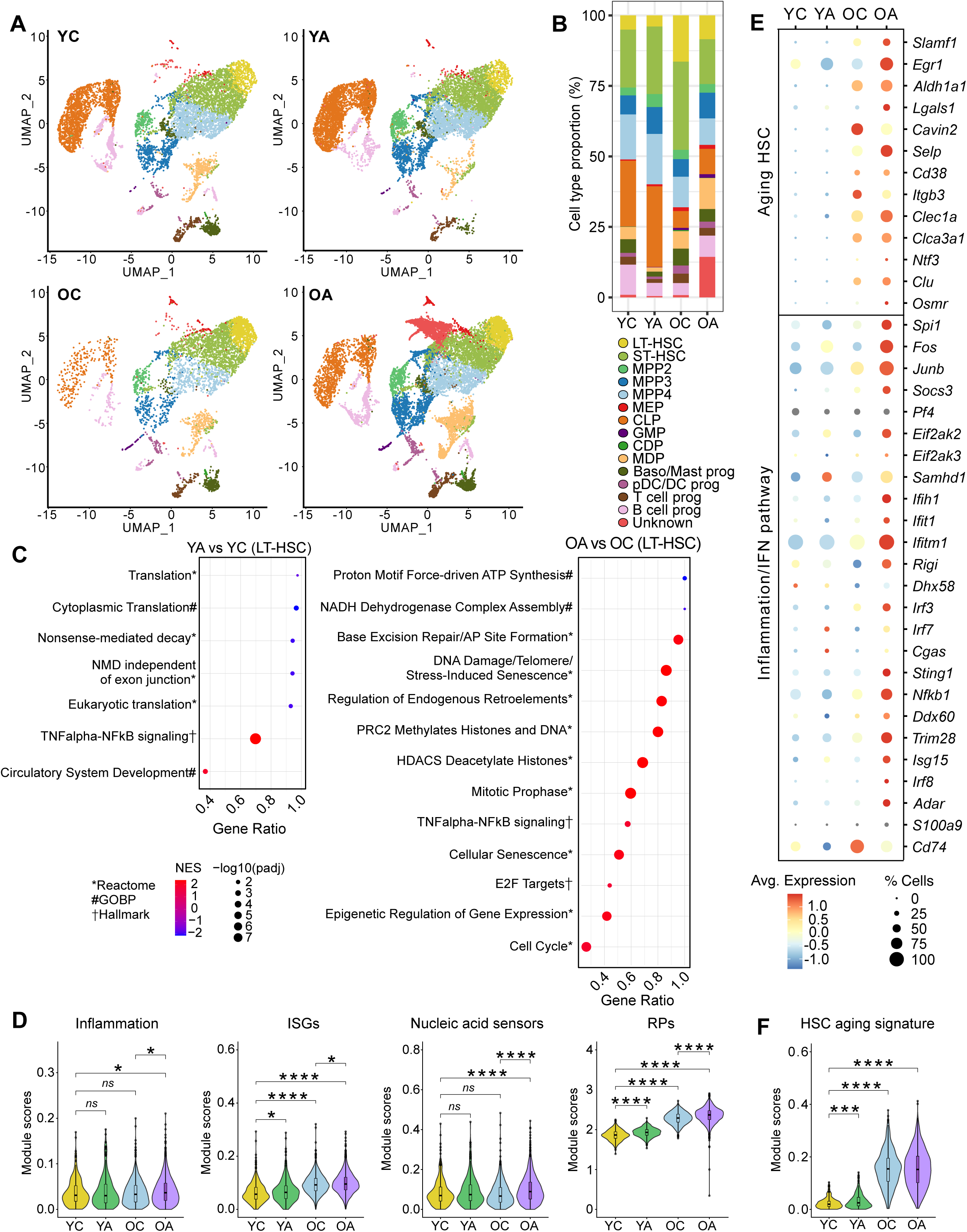
Chronic alcohol consumption promotes inflammation and alters the age-associated transcriptional program in aged LT-HSCs. (A) scRNA-seq UMAP embedding of flow-sorted murine LSK cells showing 4 groups. (B) Proportion of cell type clusters in YC, YA, OC, and OA groups. (C) Top gene set enrichment analysis (GSEA) pathways enriched in LT-HSCs of YA and OA mice as compared to their respective age-matched control. GSEA was calculated from differentially expressed genes that passed a p < 0.01 cutoff, which were subsequently ranked based on average Log2FC before calculation using fgsea package. The whole list is in **Table S5-6**. (D) Calculated module scores for genes related to inflammation, interferon-stimulated genes (ISGs), nucleic acid sensors, and RPs comparing 4 groups. (E) A dotplot showing normalized expression of selected HSC aging, inflammation, and interferon pathway genes in 4 groups. (F) HSC aging signature score calculated based on genes published in Svendsen et al, 2021^44^. scRNA-seq, single-cell RNA sequencing; log2FC, log2 fold change; RP, ribosomal protein.

Differential expression analysis revealed that chronic alcohol consumption altered 123 and 497 genes in young and aged LT-HSCs based on |log2FC| ≥ 0.58 & FDR ≤ 0.05, respectively (**Figure S3C**, **Table S1**). In contrast, aging alone (OC vs YC) led to 1937 DEGs (**Figure S3C**, **Table S1**). GSEA of young LT-HSCs revealed that chronic alcohol consumption upregulated TNFα-NFκB signaling and downregulated nonsense-mediated decay and protein translation pathways (**Figure 3C, Table S2**). In contrast, in addition to TNFα-NFκB signaling, aged LT-HSCs upregulated base excision repair (BER)/Apurinic/apyrimidinic (AP) site formation, DNA damage responses, senescence, and cell cycle programs, while downregulating pathways involved in proton motive force-driven ATP synthesis and NADH dehydrogenase complex activity following alcohol drinking (**Figure 3C, Table S2**). Furthermore, chronic alcohol consumption upregulated pathways related to epigenetic regulation in aged LT-HSCs, including nucleosome reorganization, histone deacetylation, chromatin modification, epigenetic regulation of gene expression, and regulation of endogenous retroelements—all of which are downregulated in physiological aging (**Figure S3D, Table S2**)^17,18^, suggesting alcohol-induced chromatin remodeling in aged LT-HSCs.

Chronic alcohol consumption upregulated inflammation, interferon-stimulated genes (ISGs), nucleic acid sensors, and ribosomal proteins (RPs) in LT-HSCs (**Figure 3D**). Several ISGs and dsRNA sensors, such as *Irf3*, *Irf8*, *Trim28*, *Rigi*, *Eif2ak2*, *Ifitm1*, and *Ifih1,* were upregulated only in aged LT-HSCs by alcohol exposure (**Figure 3E**). Notably, chronic alcohol consumption upregulated HSC aging genes^44^, including *Selp*, *Lgals1*, *Clec1a*, *Osmr*, and *Clu* (**Figure 3E**). Alcohol exposure also increased the HSC aging signature score, more significantly in young LT-HSCs (**Figure 3F**). Furthermore, upstream progenitors in old mice exhibited the highest module scores for inflammation, ISGs, nucleic acid sensors, and RPs due to alcohol (**Figure S4A-D**). p53 activity, as measured by the Haem p53 module score^45^, was at the highest level in alcohol-exposed aged LT-HSCs, correlating with the expression of *Cdkn1a* and *Gadd45a* (**Figure S4E-H**). In summary, aging and chronic alcohol consumption acted synergistically to drive transcriptional changes involved in DNA damage repair, epigenetic regulation, cellular aging, and inflammation, leading to alcohol-induced accelerated inflammaging in HSCs.

### Chronic alcohol consumption promotes myeloid differentiation transcriptional programs in HSPCs

To examine whether chronic alcohol consumption promotes myeloid-biased transcriptional programs, we calculated neutrophil and monocyte differentiation scores projected onto a UMAP (**Figure 4A**). Interestingly, both module scores were significantly elevated in HSC (LT- and ST-HSC) and MPP (MPP 2/3/4) clusters (**Figure 4A-B**), suggesting that chronic alcohol consumption induces myeloid-biased transcriptional programs from the most undifferentiated HSCs. Several TFs, such as *Spi1* (PU.1), *Cebpa* (CEBPα), *Cebpd* (CEBPδ), and *Irf8* (IRF8), are crucial in driving the differentiation of HSC into myeloid cells^46–48^. Importantly, cytokine receptors such as colony-stimulating factor 1 and 2 receptors (CSF1R and CSF2R) dictate lineage commitment to monocytes and granulocytes^49,50^. In line with increased neutrophil and monocyte differentiation module scores, we found increased expression of genes encoding PU.1, CEBPα, CEBPδ, IRF8, as well as CSF1R and CSF2R in both HSC and MPP clusters following chronic alcohol consumption (**Figure 4C-F**), with the highest expression levels observed in OA mice. Taken together, our data suggest upregulation of the myeloid transcriptional programs in HSCs and MPPs due to chronic alcohol consumption, which was perpetuated with aging.

**Figure 4.**
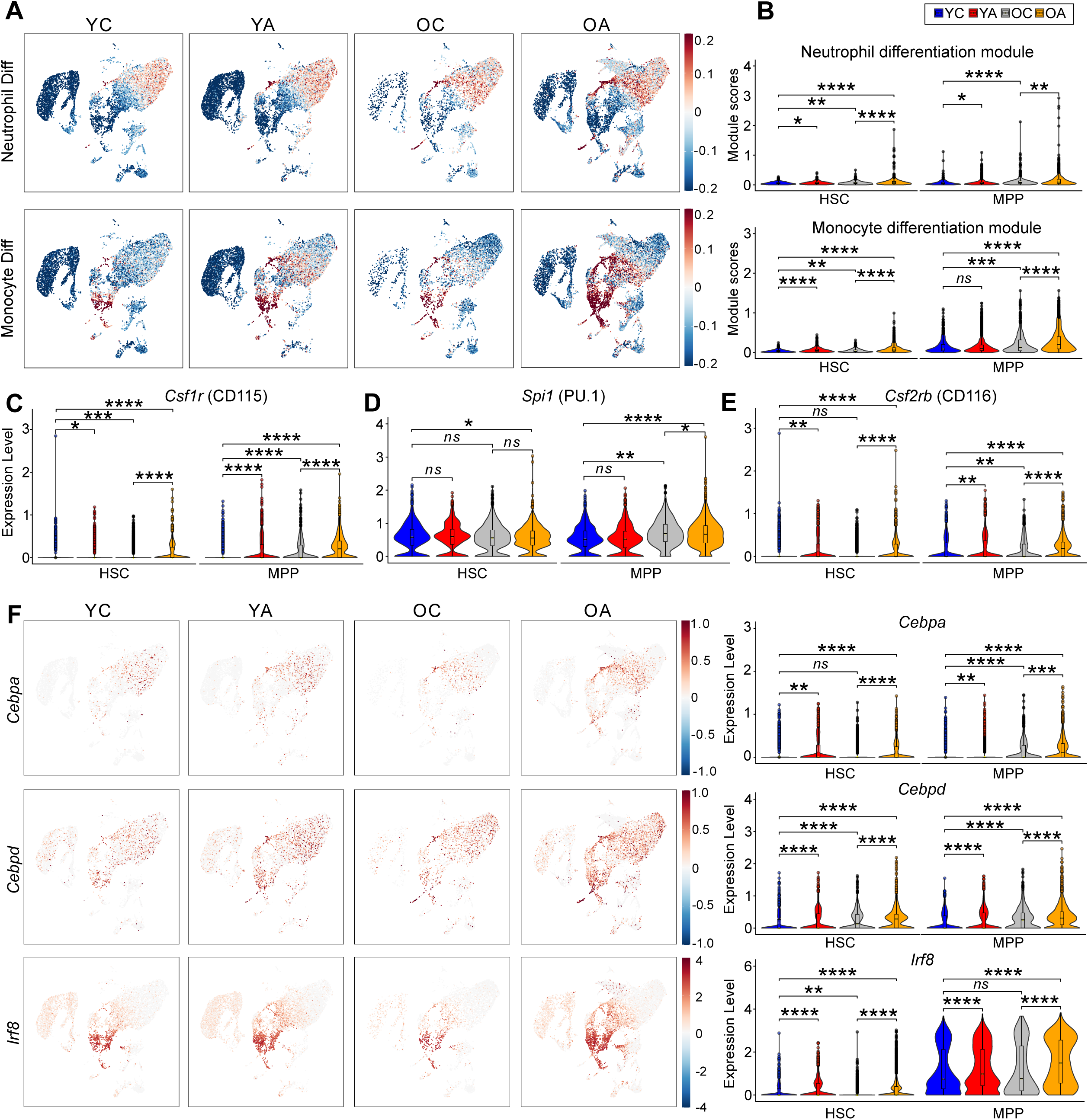
Chronic alcohol consumption upregulates the myeloid transcriptional programs in HSPCs. **(A)** UMAP-projected neutrophil and monocyte differentiation module scores. **(B)** Quantification of neutrophil and monocyte module scores in HSC (combined LT-HSC and ST-HSC) and MPP clusters (combined MPP2, MPP3, MPP4). **(C-E)** Violin plots showing normalized expression of *Csf1r* **(C)**, *Spi1* **(D)**, and *Csf2rb* **(E)** genes in HSC and MPP clusters. **(F)** UMAP projection of myeloid TFs *Cebpa*, *Cebpd*, and *Irf8,* as well as calculated module score in HSCs and MPPs.

### Chronic alcohol consumption perpetuates TE upregulation in aged murine LT-HSCs

Alterations in epigenetic and nucleic acid-sensing pathways (**Figure 3C-E**) suggested a possibility of alcohol-induced epigenetic TE derepression. To test this, the scTE pipeline was used to incorporate TE in scRNA-seq analysis^51^. Chronic alcohol consumption induced more TE upregulation in aged than in young LT-HSCs (**Figure 5A, S5A, Table S3**). LTRs constituted the major TE class showing the highest expression level among differentially expressed (DE) TE in both age groups (**Figure 5A-B**). Unlike the YA vs. YC comparison, LINEs and SINEs were also mostly upregulated in OA vs. OC, with the highest expression of these classes found in the OA group (**Figure 5A, S5A**). For example, LTR species MLTR31C-Mm and the intracisternal A particle IAPEz-int were upregulated in both young and old alcohol-treated LT-HSCs, whereas LTR75-1 and MMTV-int were upregulated only in alcohol-treated old LT-HSCs (**Figure 5B, S5A; Table S3**).

**Figure 5.**
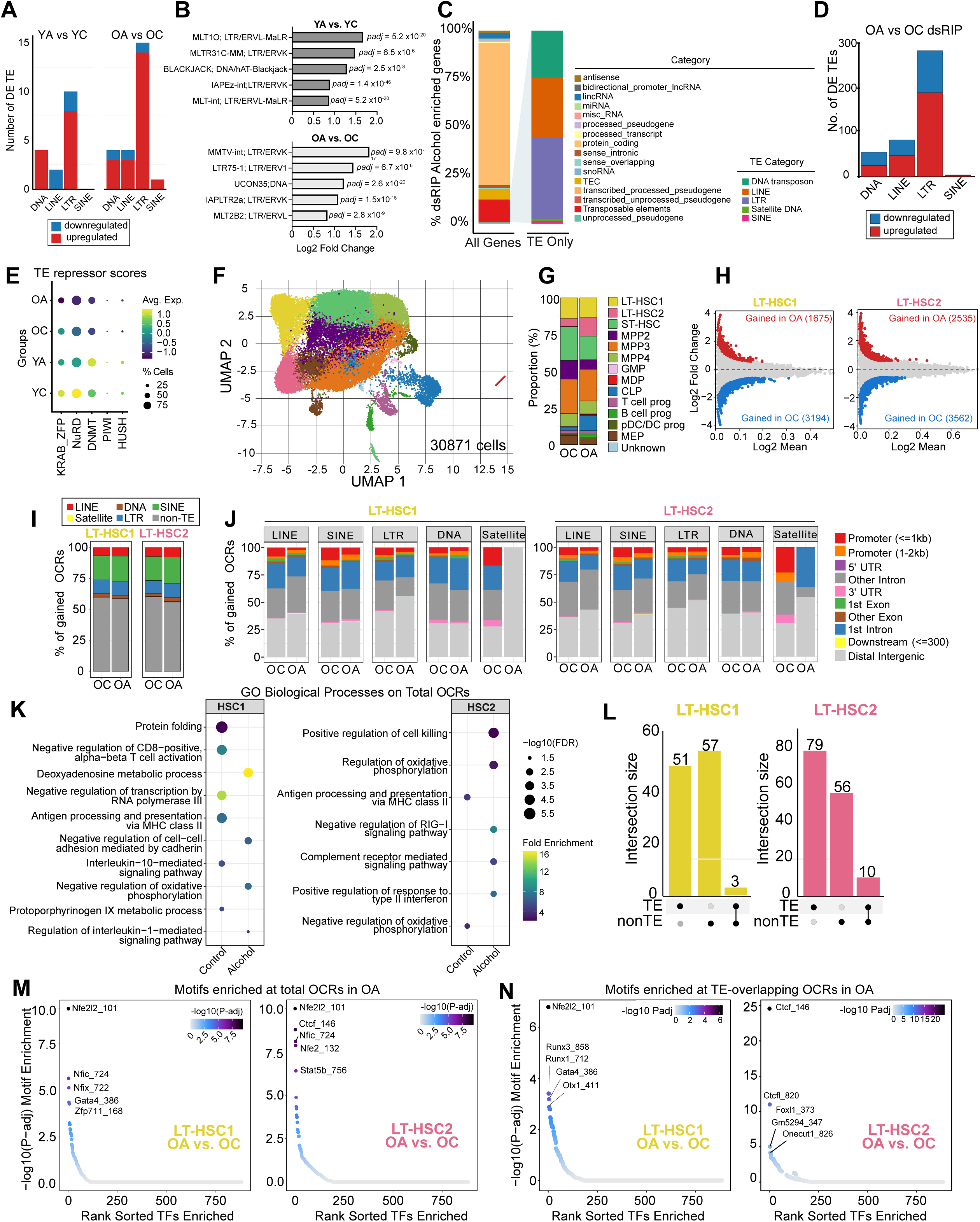
Chronic alcohol consumption and age synergistically derepress TE in HSPCs. (A) Stacked barplots showing numbers of differentially upregulated and downregulated TEs in YA vs. YC and OA vs. OC scRNA-seq. Number of DE TEs was based on padj ≤ 0.05 and absolute [log2FC] ≥ 0.58. (B) Top 5 significantly upregulated TEs in YA vs. YC and OA vs. OC LT-HSC based on Log2FC with indicated *padj* value. (C) Distribution of dsRNAome (dsRNA species) with respect to genomic annotations and TE classes in OA vs. OC total bone marrow cells (n = 5 mice/group). (D) Stacked barplots showing numbers of differentially enriched TEs within OA vs. OC enriched dsRNAs in mouse bone marrow cells enumerated based on padj ≤ 0.01 and absolute [log2FC] > 0.1 (n = 5 mice/group). (E) A dotplot of LSK scRNA-seq showing calculated module scores for several TE repressor gene families across four conditions. (F) scATAC-seq UMAP showing global open chromatin profile of LSK cells. Cell type was identified by projecting scRNA-seq clusters into scATAC-seq UMAP by using addGeneIntegrationMatrix() function and confirmed with gene accessibility of bona-fide HSPC. Clusters are color-coded to the same scRNA-seq clusters, except for the LT-HSC1 and LT-HSC2 subclusters. (G) Cell type proportion from clusters shown in scATAC-seq **UMAP**. (H) MA plot showing gained OCRs in OA vs. OC comparison. Gain in OA (red) and gained in OC (blue) OCRs were filtered based on FDR ≤ 0.05 and |Log2FC| ≥ 0.58. (I) Bar plots showing the distribution of gained OCRs overlapping with TE classes in LT-HSC1 and LT-HSC2 of OA and OC groups. (J) Bar plots showing the distribution of TE-overlapping OCRs based on genomic locations in LT-HSC1 and LT-HSC2 of OA and OC groups. (K) Dot plot showing selected GO biological processes enriched in total OCRs of control and alcohol groups. Dot size represents -log10 FDR, color gradation represents Fold Enrichment. The whole list is in Supplemental **Table S5**. (L) UpSet plots showing the number of significantly enriched TF binding motifs in TE- or non-TE-overlapping OCRs found in the OA group. Set size indicates motifs that passed FDR ≤ 0.05 and log2FC ≥ 0.58. The full list of TF motifs is in Table S7 (**M-N**) Motif enrichment plots showing the top 5 rank-sorted TF binding motifs from total (**L**) and TE-overlapping (**M**) OCRs enriched in LT-HSC1 and LT-HSC2 of the OA group. The whole lists is in Supplemental **Table S8**. **TE**, transposable element; **LTR**, long terminal repeats; **LINE**, long interspersed nuclear element; **SINE**, small interspersed nuclear element, **FACS**, fluorescence-activated cell sorting; **scATAC-seq**, single cell assay for transposase-accessible chromatin sequencing; **FDR**, false discovery rate; TF, transcription factor; CIS-BP, catalog of inferred sequence binding preferences; **MA plot**, M (log ratio) vs. A (average) plot; **OCRs**, open chromatin regions.

Since TEs can form dsRNA^52,53^, we asked whether chronic alcohol consumption induced bone marrow cells to accumulate TE-derived dsRNAs *in vivo*. To this end, we performed dsRNA immunoprecipitation using the J2 antibody (J2-RNA-Immunoprecipitation; J2-RIP) from total RNA harvested from control- and alcohol-fed mouse bone marrow cells. We sequenced both J2-RIP and INPUT samples harvested from OC and OA mice. The amount of IgG-immunoprecipitated RNA was insufficient for library preparation, confirming the J2 antibody’s specificity. We identified 601 genes enriched in OA J2-RIP compared to OC J2-RIP, after normalization for INPUT variations (**Figure S5B, Table S4**). In line with the previous report in yeast^54^, protein-coding genes constituted the major components of dsRNAs (**Figure 5C**). Interestingly, TEs accounted for the second-largest component of dsRNAs, comprising up to 15% of the total OA-enriched J2-RIP genes. Processed pseudogenes comprised the third major dsRNAome enriched in OA, followed by long non-coding RNAs. Mirroring differentially-expressed (DE) TE abundance in scRNA-seq data, LTRs were dominantly enriched in OA J2-RIP, followed by LINEs and DNA transposons, whereas SINEs and other repetitive elements were minor (**Figure 5C-D**). LTR species, such as MER34A1 and RMER19B2, and LINE species, such as L1M3de, were enriched in dsRNA (**Figure S5C**). Surprisingly, we did not detect mitochondrial-derived dsRNAs in our dsRIP-seq dataset, unlike previously reported^55,56^. Overall, these experiments suggest that chronic alcohol consumption upregulates TEs, with the strongest response observed in aged HSCs, thereby contributing to dynamic changes in the cells’ dsRNAome.

### Chronic alcohol consumption increases accessibility of transposable elements in aged HSCs

We observed that TE repressors wane with aging^26,57–59^ and alcohol exposure (**Figure 5E**), which led us to hypothesize that alcohol exposure alters chromatin accessibility of TEs in HSCs. To prove this, we performed a single-cell assay for transposase-accessible chromatin with sequencing (scATAC-seq) on LSK cells from OA and OC mice. Surprisingly, we identified two LT-HSC clusters, LT-HSC1 and LT-HSC2, based on chromatin accessibility of bona fide LT-HSC marker genes (**Figure 5F, S6A-B**). LT-HSC1 showed greater accessibility for *Slamf1* and *Mecom*, whereas LT-HSC2 exhibited greater accessibility for *Procr, Pdzk1ip1*, *Mpl*, and *Hlf*. *Aurkb* was closed in LT-HSC1 but open in LT-HSC2, suggesting a more active cell cycle in LT-HSC2, given Aurora kinase’s role in mitosis^60^ (**Figure S6B**). From 158,707 cluster-specific open chromatin regions (OCRs) (**Figure S6C**), several transcription factor (TF) motifs, including FOS, SMARCC1, BACH1, and BACH2 TF, were shared by LT-HSC1 and LT-HSC2, whereas the CTCF motif was exclusively enriched in LT-HSC2 (**Figure S6D**). Furthermore, alcohol exposure caused LT-HSC2 expansion but reduced ST-HSC and MPP4 populations, while the LT-HSC1 remained unchanged (**Figure 5F-G**).

Differential accessibility analysis comparing OA vs. OC revealed that 1,675 and 3,194 OCRs were gained in OA and OC LT-HSC1, respectively, and 2,535 and 3,562 OCRs were gained in OA and OC LT-HSC2, respectively (**Figure 5H**). Surprisingly, alcohol exposure increased the number of OCRs overlapping with TEs, especially in LT-HSC2 (**Figure 5I**). SINE contributed to the most overlapping TE class, followed by LTR and LINE classes. To find patterns of genomic distribution of TE-overlapping OCRs, we calculated the proportion of TE-overlapping OCRs located within various genomic locations. For almost all TE classes, except for DNA transposons, we found a pattern in which alcohol reduced TE-overlapping OCRs in promoters and exons, whereas it increased the proportion of TE-overlapping OCRs in intergenic and intronic regions (**Figure 5J**). For example, we found Alu:SINE:B1- and SINE:ID4-overlapping OCRs at intergenic regions nearby *Cebpa* (**Figure S7A**).

GO analysis of genes associated with total and TE-overlapping OCRs revealed enrichment of inflammatory and interferon pathways in OA LT-HSC1 and LT-HSC2, whereas MHC class II and antigen-presentation pathways were enriched in OC LT-HSC1 and LT-HSC2 (**Figure 5J, S7B, Table S5-6**). Notably, complement receptor-mediated signaling and negative regulation of HSC differentiation pathways were specifically enriched in TE-overlapping OCRs in OA LT-HSC2, whereas the retrotransposon silencing pathway was only enriched in OC LT-HSC2 (**Figure S7B, Table S6**). In line with reduced erythrocyte numbers and hemoglobin levels in alcohol-exposed old mice (**Figure S1E, Table S5-6**), pathways responsible for erythrocyte differentiation, hemoglobin biosynthesis, and protoporphyrinogen IX metabolic processes were enriched only in control aged LT-HSCs (**Figure 5K, S7B, Table S5-6**).

TEs can act as *cis*-regulatory elements providing TF binding sites^31,61^. LT-HSC1 exhibited comparable motif enrichment in both TE and non-TE-overlapping OCRs, whereas LT-HSC2 showed dominant enrichment for TF motifs found in TE-overlapping OCRs (**Figure 5L, Table S7**). Motif enrichment analysis in total OCRs revealed NFE2L2 and NFIC motifs enrichment in both LT-HSCs, whereas GATA4 and ZFP711 were found exclusively in LT-HSC1, and CTCF and STAT5b motifs were found exclusively in LT-HSC2 (**Figure 5M**). Interestingly, motif enrichment analysis of TE-overlapping OCRs revealed greater specificity and exclusivity between HSC clusters. For example, NFE2L2, RUNX3, RUNX1, and GATA4 were among the top motifs in LT-HSC1, whereas CTCF, ONECUT1, and FOXL1 motifs were enriched in TE-overlapping OCRs in LT-HSC2 (**Figure 5N**). Furthermore, chromVAR deviation analysis unraveled that CTCF motifs exhibited the highest mean difference of Z-score between OA vs. OC in LT-HSC2 (**Figure S7C**). Taken together, our results suggest that chronic alcohol consumption increases chromatin accessibility at TE-harboring distal intergenic regions.

### Chronic alcohol consumption promotes myeloid bias and inflammation in xenotransplanted human HSPCs

To investigate the effects of chronic alcohol consumption on human HSPCs, we employed a xenotransplantation model, in which healthy human BM CD34^+^ cells were transplanted into immunocompromised NBSGW mice (**Figure 6A**). We performed two independent cohort experiments from different donors. Cohort 1 was from a 27-year-old female, and cohort 2 was from a 24-year-old male. Following confirmation of human cell engraftment at 6 weeks post-xenotransplantation (**Figure 6B**), mice were fed with either a liquid diet containing 5% (v/v) alcohol or an isocaloric control liquid diet for 8 weeks. We achieved a mean BEC of approximately 40 mM in the alcohol group, and no significant differences in diet intake or body weight changes between the groups (**Figure S8A-B**), and comparable human cell engraftment in PB and BM in both groups following alcohol exposure (**Figure 6C**). However, there was a significant increase in CD14^+^ monocytes, CD15^+^ granulocytes, and CD33^+^ myeloid cells, whereas the CD19^+^ B cell proportion decreased, resulting in a higher myeloid to lymphoid (M:L) ratio in the alcohol group (**Figure 6D**). Notably, human T cell engraftment was not observed, presumably due to thymic atrophy in NBSGW mice^62^. Overall, HSPC frequencies did not change, but a slight increase in MPP was observed in the alcohol group (**Figure S8C**). To assess the effects of chronic alcohol exposure on long-term engraftment potential, we performed secondary transplantation by transplanting 500,000 sorted human CD45+CD34 cells (cohort 1) or 10 million unsorted bone marrow cells (cohort 2) from the alcohol feeding experiments into NBSGW mice without conditioning. Secondary transplant recipients were monitored up to 16 weeks without further alcohol treatment. We observed slightly lower human engraftment levels in previously alcohol-exposed groups; however, the differences did not reach statistical significance (**Figure 6E**).

**Figure 6.**
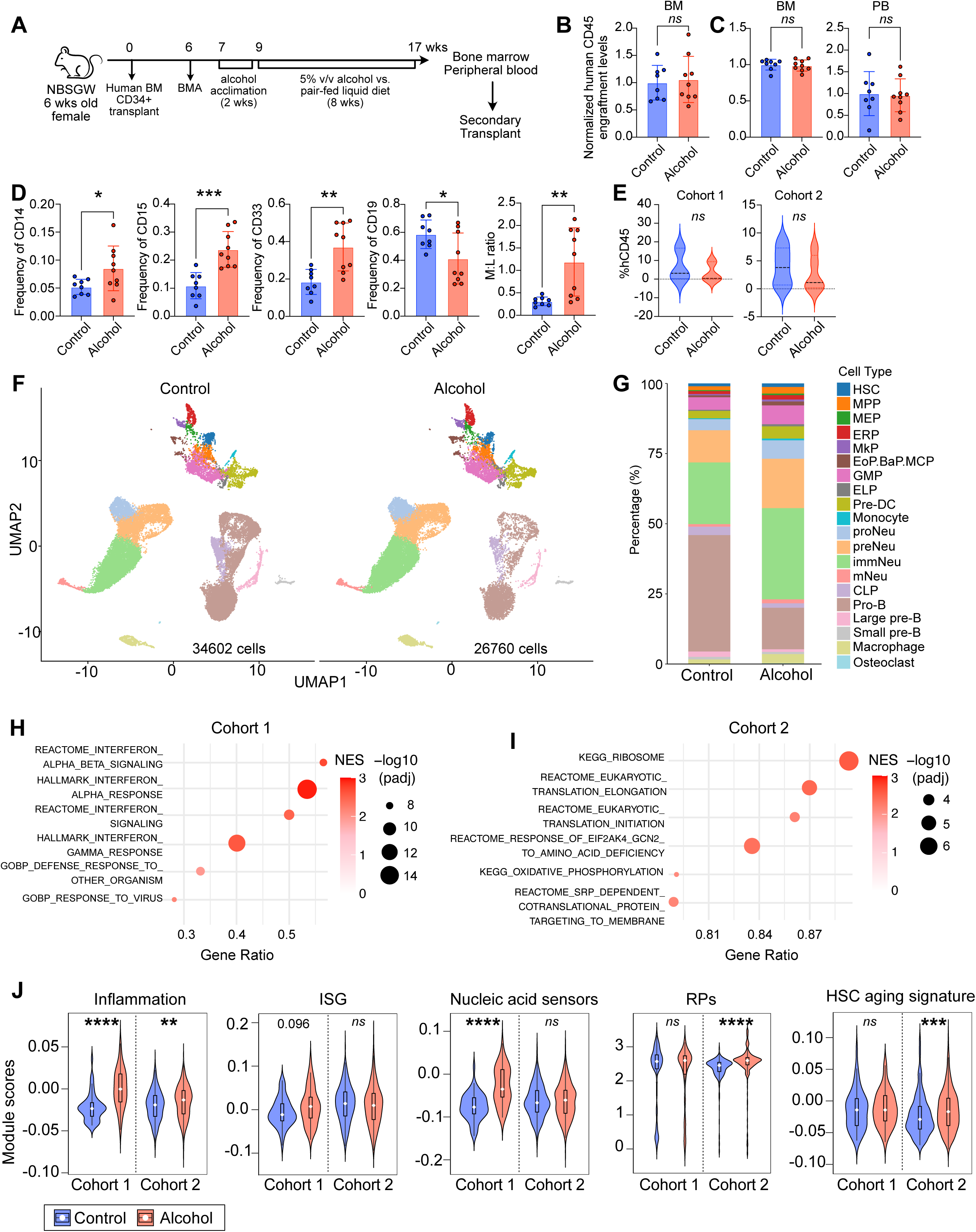
Chronic alcohol consumption caused myeloid bias and inflammation in xenotransplanted human HSPCs. **(A)** Experimental scheme of human HSPC xenotransplantation followed by alcohol feeding experiment. Two independent experiments were performed. Cohort 1 was from a 27-year-old female donor, and cohort 2 was from a 24-year-old male donor. **(B)** Human CD45+ cell engraftment before chronic alcohol feeding (baseline) **(C)** Human CD45+ cell engraftment level after chronic alcohol feeding (endpoint). **(D)** Proportion of human monocytes (CD14+), granulocytes (CD15+), myeloid cells (CD33+), B cells (CD19+), and ratio of myeloid vs. lymphoid cells in peripheral blood. **(E)** Human CD45+ engraftment levels at 16 weeks post-secondary transplant are shown for each cohort (cohort 1: n=3 per group; cohort 2: n=5 per group). **(F)** UMAP of sorted human CD34+CD45+ stem progenitor cells obtained at the endpoint of the alcohol feeding experiment. Cell clusters were color-coded. **(G)** Cell-type proportions showing the distribution of each cluster in the alcohol and control groups. **(H-I)** GSEA showing the top upregulated pathways in human xenotransplanted HSCs following chronic alcohol exposure in cohort 1 (**H**) and cohort 2 (**I**). **(J)** Violin plots showing module score calculation of inflammation, ISG, nucleic acid sensors, RPs, and HSC aging signature genes. **(B-D)** All data were pooled from 2 independent experiments and analyzed using a Student’s t-test. *p < 0.05; **p < 0.01; ***p < 0.001; ****p < 0.0001. Bars indicate mean ± SD, dots represent individual mice. **(F-G)** Pooled data shown from 2 independent experiments (cohort 1: n=1 per group; cohort 2: n=2 per group; combined cohort: n=3 per group). **HSPC**, hematopoietic stem progenitor cells; **UMAP**, Uniform Manifold Approximation and Projection; **GSEA**, gene set enrichment analysis; **IFN**, interferon; dsRNA, double-stranded RNA; **ISG**, interferon-stimulated genes; **RPs**, ribosomal proteins.

Single-cell RNA-seq (scRNA-seq) showed 20 UMAP clusters of human CD45+ CD34+ cells following unsupervised clustering, including HSCs, MPPs, megakaryocyte-erythroid progenitors (MEPs), and downstream progenitors (**Figure 6F**). Consistent with the PB flow cytometry data, our scRNA-seq data showed a myeloid bias, with the increased and decreased numbers of neutrophil progenitors and pro-B cell clusters, respectively, in both cohorts (**Figure 6G, S8D-E**; results from combined cohorts shown). Interestingly, each cohort exhibited distinct molecular perturbations following alcohol exposure that converged into myeloid bias and heightened inflammation. Specifically, cohort 1 HSC cluster significantly upregulated interferon pathways (**Figure 6H**), whereas cohort 2 HSC cluster mainly upregulated ribosome and eukaryotic translation elongation pathways (**Figure 6I**). Cohort 1 was further characterized by a significantly higher nucleic acid sensor module score and a trend toward increased interferon-stimulated genes (ISGs: *IFITM1*, *IFITM3*, *IFI6*, and *ISG15*) module score, suggesting heightened interferon responses caused by nucleic acid sensor activation (**Figure 6J**). Conversely, cohort 2 was characterized by significantly higher ribosomal protein module score and HSC aging signature, suggesting perturbed proteostasis and accelerated aging (**Figure 6J**). These data suggest significant inter-individual heterogeneity in HSCs’ molecular response to chronic alcohol exposure, culminating in myeloid bias and heightened inflammation.

### Chronic alcohol exposure also upregulates TEs in human HSPCs

To validate alcohol-induced TE upregulation in human HSPCs, we analyzed the scRNAseq data using the scTE pipeline. We found cohort-specific TE upregulation in HSC (**Figure 7A**) and MPP (**Figure 7B**) populations, with ERV1 upregulation being shared by both cohorts’ HSC and MPPs. ERV1 upregulation was also most prominent in TE analysis of bulk RNA-sequencing data of *in vitro* alcohol-treated human BM CD34+ cells (**Figure 7C**), which also showed higher dsRNA staining with J2 monoclonal antibody (**Figure 7D**). We confirmed the significant upregulation of IFNA1 and IFNB1, but not IFNG by RT-qPCR of *in vitro* alcohol-treated human CD34+ BM cells. HeLa cells, as well as *in vivo* alcohol-exposed mouse alveolar macrophages showed intensified dsRNA staining (**S8F**). Furthermore, human leukemia cell line MOLM-13 showed elevated dsRNA concentration following alcohol treatment *in vitro* (**Figure S8G**). Overall, our data suggest that chronic alcohol consumption upregulates TEs, resulting in dsRNA accumulation in the cytosol, which activates dsRNA sensors and elicits interferon responses in human HSPCs.

**Figure 7.**
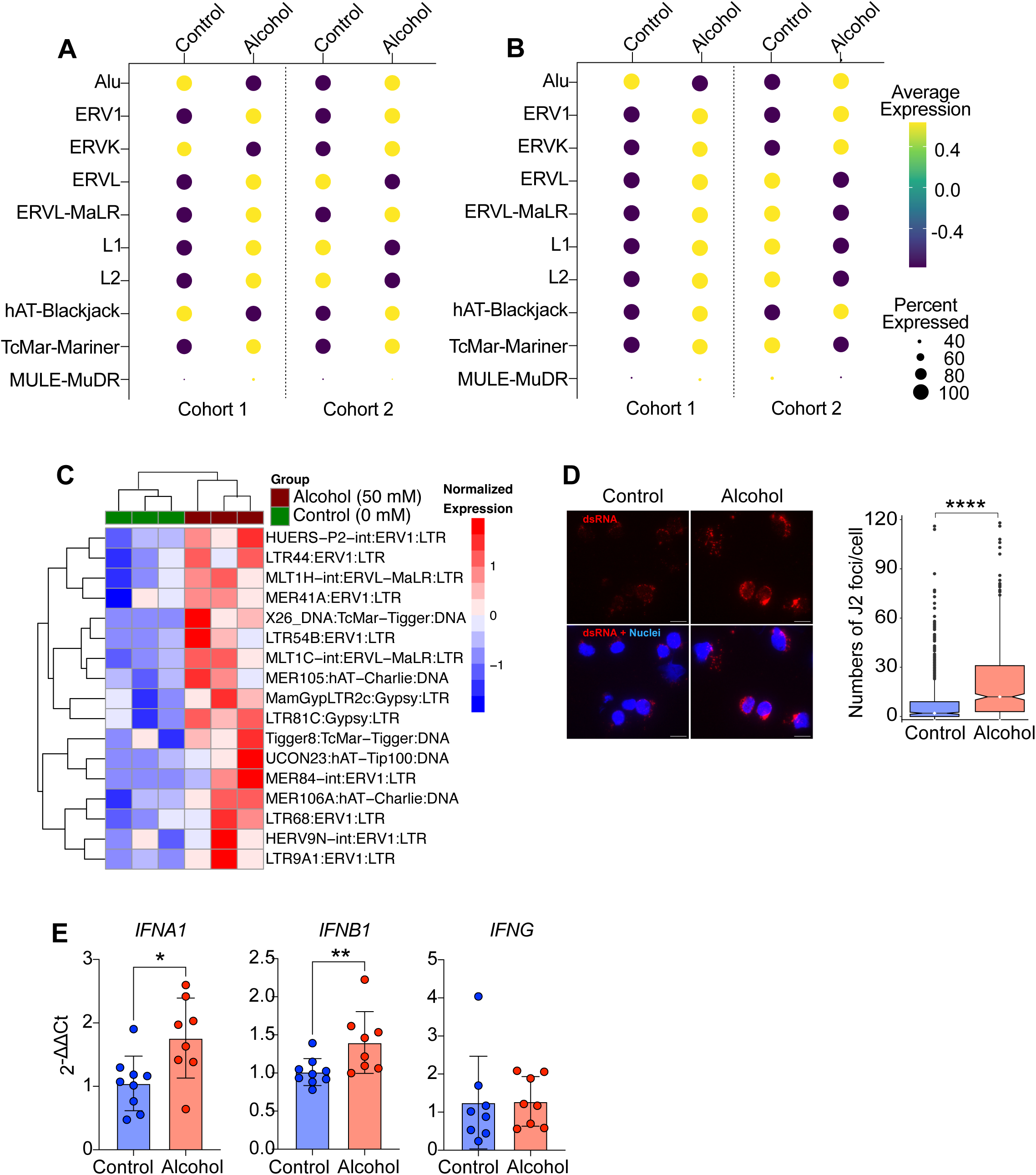
Alcohol exposure led to upregulation of transposable elements and dsRNA accumulation in human HSPCs. **(A-B)** scTE analyses of HSCs **(A)** and MPPs **(B)** are shown for each cohort. **(C)** TE analysis of bulk RNA-seq from *in vitro* alcohol-treated HSPCs shows upregulation of LTRs, most notably ERV1. Human BM CD34+ cells were treated with alcohol at 0 vs. 50 mM for 48 hours in three biological replicates in one experiment. **(D)** Immunofluorescence staining of dsRNAs in alcohol-exposed cultured human BM CD34+ cells. Quantification of dsRNA foci/cell was performed using an automated pipeline in CellProfiler software (v.4.2.6). **(E)** RT-qPCR for IFNA1, IFNB1, and IFNG. Human BM CD34+ cells were treated with alcohol at 0 vs. 50 mM for 48 hours in four biological replicates in two experiments. **dsRNA**, double-stranded RNA; **scTE**, single-cell Transposable Elements; **HSCs**, hematopoietic stem cells; **MPPs**, multipotent progenitor populations; **LTR**, long terminal repeats; **ERV1**, endogenous retrovirus 1; **RT-qPCR**, reverse-transcription quantitative PCR.

## Discussion

Chronic alcohol consumption has been shown to cause inflammation via increased differentiation of inflammatory monocytes and osteoclasts in non-human primate models^14,15^. Here, we report previously unrecognized cellular and molecular consequences of chronic alcohol consumption on murine and human HSPCs, which are strongly modulated by age in mice and by donor variability in humans. We achieved a mean BEC of approximately 40 mM, which is commonly observed in individuals with heavy drinking. Young LT-HSCs demonstrated minimal myeloid bias and limited transcriptional changes. By contrast, aged LT-HSCs displayed a pronounced myeloid bias and extensive transcriptomic changes in response to alcohol exposure, including DNA damage responses, cell-cycle regulation, senescence, histone modification, endogenous retroelement control, and inflammation. Notably, chronic alcohol exposure upregulated TEs in both age groups, with the OA group showing more intense TE expression, associated with increased chromatin accessibility at TE loci in intronic and distal intergenic regions. In xenotransplanted human HSPCs, alcohol exposure also induced myeloid bias, type I interferon signaling, upregulation of dsRNA sensors, and TE reactivation from ERV families. Moreover, the upregulation of RP genes was also conserved across species. These molecular alterations are likely to correspond to the LT-HSC2 subset on scATAC-seq, which is expanded in response to alcohol exposure and whose epigenomic features, including CTCF motif enrichment, resemble those of inflammatory memory HSCs reported in a recent preprint^22^ or previously described activated HSCs^63^. Despite these perturbations, to our surprise, human and murine HSPCs retained long-term repopulation capacity. However, alcohol-exposed HSPCs exhibited a persistent myeloid bias and reduced capacity to enter the cell cycle following a secondary inflammatory challenge, even after cessation of alcohol exposure, suggesting trained immunity^47,64^ and aging^41,42^ in HSCs, respectively.

Previous studies highlight the role of TE upregulation during HSC emergence^65^, demand-induced erythropoiesis (e.g., during pregnancy and blood loss)^66^, and chemotherapy-induced stress hematopoiesis^67^. These studies suggest that TEs, along with cellular machineries that sense them, play a critical role in initiating HSC differentiation toward myeloid and erythroid lineages. This process may be partly driven by sensing of TE-derived DNA via the cGAS-STING pathway^57,66^ and TE-derived dsRNA via dsRNA-sensing pathways (e.g., MDA5, RIG-I, PKR, OAS, etc.)^67^, ultimately leading to activation of interferon signaling and inflammatory responses^57,67^. Consistent with these findings, we observed that chronic alcohol consumption activated interferon and inflammatory pathways and upregulated dsRNA sensor genes in both human and murine HSCs. Given the known role of MDA5 in recognizing TE-derived dsRNA^67^ and aging via perturbed proteostasis^68^, our findings of upregulation of dsRNA sensors and RPs in alcohol-exposed aged HSCs suggest that similar principles may operate in alcohol-induced inflammaging. Additionally, we detected an accumulation of TE-derived dsRNAs, which represent the second most abundant class of dsRNA species upon alcohol exposure in the BM of old mice. This observation aligns with previous reports describing the accumulation of TE-derived dsRNAs in response to cellular stress^52^.

Furthermore, TEs harbor TF-binding sites that can contribute to gene regulatory programs^69^. Enrichment of the NFE2L2 motif inside TE-overlapping OCRs suggests a potential role for TEs in regulating oxidative stress response^70^ and inflammatory signaling^32^. Notably, the presence of the CTCF motif within TE-overlapping OCRs raises the possibility that TEs may also participate in genome structural regulation^31^, potentially mediating chronic alcohol-induced inflammation in aged LT-HSCs. Establishing how TE mediates alcohol-induced inflammation warrants future studies.

Besides TE upregulation, our work showed that chronic alcohol consumption increased cell cycle entry (Ki67) and γ-H2AX levels in LT-HSCs, accompanied by increased p53 activity score and RP transcript accumulation, indicative of replicative stress^18,71^ and genotoxic stress^72^, which are characteristics of HSC aging. We identified a novel synergistic effect of aging and chronic alcohol consumption in amplifying HSC aging phenotypes, including induction of DNA damage responses, cellular senescence programs, dsRNA-sensing transcriptional programs, inflammation, and myeloid bias in aged HSCs. Although these alterations did not significantly increase the HSC aging module score, we found transcript levels of genes implicated in HSC aging^44^, such as *Selp*, *Lgals1*, *Clec1a*, *Osmr*, and *Clu,* were significantly upregulated in alcohol-exposed aged LT-HSCs.

Surprisingly, chronic alcohol exposure led to the contraction of the aged LT-HSC compartment, which was the opposite of the age-induced HSC expansion. This shrinking of the aged LT-HSC compartment under alcohol exposure could be explained by the increased cell cycle reflecting precocious myeloid differentiation during IL-1-driven myelopoiesis^73^, or due to alcohol-induced LT-HSC deaths as previously reported in interferon-induced HSC apoptosis^74^.

We discovered a significantly increased vulnerability of aged HSCs to alcohol exposure. First, we observed that aged HSCs exhibited a significant reduction in epigenetic repressors compared with young HSCs, including KRAB-ZFPs, PIWI, NuRD, and DNMTs known to silence TEs^24–26,52,76^. The reduction in epigenetic silencing capacity, which was further decreased by alcohol exposure, may have compromised epigenome maintenance and potentially triggered widespread epigenomic alterations^17,18,77^, leading to increased chromatin accessibility at TE-containing loci^78^. Second, aged HSCs have a reduced capacity to repair oxidative DNA damage compared to young HSCs^18,71^, leading to persistent DNA damage that triggers epigenome rearrangements^79^. Indeed, oxidative DNA damage preferentially arises in intergenic regions containing repetitive elements^80^, potentially explaining the preference for chromatin reopening at those regions.

In summary, our study reports cellular and molecular alterations in murine and human HSPCs following chronic alcohol exposure, illuminating potential interactions between alcohol consumption and aging that reinforce pathways involved in inflammation, ribosome biogenesis, epigenome remodeling, and TE dysregulation.

## Limitations

Our study has some limitations. We used xenotransplantation of human HSPCs into NBSGW mice, which do not fully support T-cell development^62^. While our data illuminate the impact of chronic alcohol drinking on HSC inflammation and TE upregulation, it remains unclear to what extent TE transposition and direct sensing promote HSC inflammaging. Our scATAC-seq was performed only in alcohol-exposed and control old mice because the most pronounced TE upregulation and transcriptomic changes were observed in this group; however, this does not exclude the possibility that alcohol exposure also alters chromatin accessibility in young mice. Lastly, it remains unclear the contribution of each HSPC subpopulation to dsRNAome, as the dsRIP experiment was conducted using total bone marrow cells to meet the dsRIP-seq input requirement. We did not test prolonged or repeated alcohol exposure, and such regimens could still compromise HSC repopulation capacity.

## Acknowledgements

This work was supported by grants from the National Institute on Alcohol Abuse and Alcoholism (R21 AA030864-01A1 – M.J.), the National Heart, Lung, and Blood Institute (R00 HL177829 – M.J), Maryland Stem Cell Research Fund (MSCRF) Launch Award (2023-MSCRFL-5999 – M.J.), Cigarette Restitution Fund Recruitment Award (PHPA-2288/M00B4600138 – M.J.), the American Society of Hematology Junior Faculty Scholar Award (M.J.), and the Maryland Stem Cell Research Fund Postdoctoral Fellowship Award (2024-MSCRFF-6407 – R.A.A.Y.). This work is partially supported by NIH S10OD026859 (Ross Flow Cytometry Core), NIH R00CA226357 (S.J.), NIH R35GM147513 (S.J.), NIH P30CA006973 (Sidney Kimmel Comprehensive Cancer Center). We thank Xiaoling Zhang, PhD, at the Ross Flow Cytometry Core, for assistance with sorting experiments for the human HSPC xenotransplantation studies. We thank Lauren K. Yum, Ph.D., Jong Seok Lee, Ph.D., Emanuele Palescandolo, Ph.D, from the Single Cell and Transcriptomics Core Facility of Johns Hopkins School of Medicine for providing assistance and service for human scRNA-seq and murine scATAC-seq experiments. We also thank Jennifer Meyers, Dixie Hoyle, and Kornel Schuebel, Ph.D., for processing our samples for the murine scRNA-seq experiment. We thank the members of the Marques laboratory (IBMC/Strasbourg, France) for providing technical expertise for dsRIP-seq. This work was carried out at the Advanced Research Computing at Hopkins (ARCH) core facility (https://rockfish.jhu.edu), which is supported by the National Science Foundation (NSF) grant number OAC1920103.

## Authorship and conflict-of-interest statements

**Contribution: R.A.A.Y** conceptualization, project administration, investigation, perform most experiments, data curation, bioinformatics analysis of aging mouse scRNA-seq, dsRIP-seq, and scATAC-seq data, formal analysis of most experiment, visualization, validation, methodology, writing-original draft, writing-review and editing, funding acquisition; **H.B** investigation, formal analysis, bioinformatics analysis of human scRNA-seq data; **V.K** investigation, formal analysis; **J.K**, **F.Y**, **H.C**, **Y.P**, **Y.F**, **Z.H**, and **J.C** investigation; **L.Z.L** helps in GSEA analysis and provide scientific input**; Z.S** resources, funding acquisition; **B.G** supervision, funding acquisition; **S.J** supervision, funding acquisition; **L.S.M.R** supervision, funding acquisition; **M.J** conceptualization, project administration, investigation, validation, supervision, writing-review and editing, funding acquisition. All authors reviewed and edited the final manuscript.

## Statement of prior presentation

Presented in poster form at the 65^th^, 66^th^ and 67^th^ annual meeting of the American Society of Hematology

## Data Sharing Statement

All sequencing data and R codes are available upon request to the corresponding author

