## Supplemental information for "Chronic alcohol exposure drives inflammaging and transposon derepression in hematopoietic stem progenitor cells"

#### Supplementary Materials and Methods

##### Mice and Husbandry

To perform human HSPC xenotransplantation experiments, 5 to 6-week-old female NBSGW mice (JAX #026622) were used. This strain allows engraftment of human HSPCs without irradiation and supports the generation of human granulocytes, monocytes, and B cells, albeit with limited robust T cell engraftment due to thymic atrophy[1]. Young (2-month-old) male and female C57BL/6NJ mice were obtained from JAX (strain #005304). Old (21 to 22-month-old) male and female C57BL/6 mice were obtained from the National Institute of Aging (NIA) Aged Rodent Colonies. For serial bone marrow transplantation experiments, female C57BL/6J (CD45.2; JAX #000664) and B6.SJL-$\mathit{Ptprc}^{a}$ $\mathit{Pepc}^{b}$/BoyJ (CD45.1; JAX #002014) were used. Mice were housed in the specific-pathogen-free animal facility in the Miller Research Building.

##### Alcohol feeding experiments

We used a modified protocol from the NIAAA alcohol feeding experiments[2]. Lieber-DeCarli isocaloric control and ethanol liquid diets were purchased from Bio-Serv (Product # F1259SP and F1258SP, respectively). Liquid diet acclimatization was performed for 1 week before the introduction of alcohol. Liquid diet intake was monitored, and access to water and chow was removed once intake stabilized after 1 week. Mice were treated with ethanol at a 1% volume/volume (v/v) dose increase every 2-3 days until reaching 5% v/v, and the treatment was continued for 4 weeks for C57BL/6 mice or 8 weeks for xenotransplanted NBSGW mice. Survival and consumption of the liquid diet were monitored daily, and body weight was monitored every 2-3 days. Of note, we reduced the alcohol dose to 4% v/v for old female mice during the 4$^{th}$ week of alcohol feeding, because we observed premature death in that group.

##### Blood ethanol concentration measurement

Approximately 50-100 μl blood withdrawn from retroorbital bleeding was quickly transferred to microcentrifuge tube and flash frozen in liquid nitrogen. Samples were then transported in dry ice and stored in -80 °C until analysis. The gas chromatography-mass spectrometry (GC-MS) was performed as previously described[3]. Briefly, 25 μl blood was aliquoted for each measurement and mixed with 5 μmol of $^{2}H_{6}$-ethanol (internal standard for ethanol) and 0.04 μmol $^{2}H_{4}$-acetaldehyde (internal standard for acetaldehyde).

Subsequently, 250 μl of 0.6 N $HClO_4$ was added to each sample. Samples were homogenized at 4 °C for 2 min using a Precellys Evolution homogenizor (Bertin), followed by centrifugation at 13,000 x g for 15 minutes at 4 °C. A 200 μl aliquot of the resulting supernatant was transferred into a 20 ml headspace vial and immediately sealed. The vials were then loaded to the GC-MS machine (Agilent Technologies) for measurement.

**Xenotransplantation of human CD34+ BM cells**

100,000 fresh (cohort 1) or 200,000 frozen (cohort 2) human CD34+ bone marrow cells were retroorbitally injected into 5 to 6-week-old female NBSGW mice without prior conditioning. Donor engraftment was validated at 6 weeks post-xenotransplantation by bone marrow aspiration (BMA) as previously described[4,5]. After confirming engraftment, the mice were assigned to either a control or an alcohol liquid diet group. After 2 weeks of alcohol acclimation, the alcohol group was fed with 5% v/v alcohol liquid diet for 8 weeks. At 17 weeks post-xenotransplantation (after 8 weeks on an alcohol- versus a control-liquid diet), mice were harvested for flow cytometry, secondary transplantation, and single-cell RNA sequencing (scRNA-seq). Secondary transplant was performed by retro-orbital injection of either 500,000 human CD34+CD45+ sorted cells (cohort 1) or 10 million unselected bone marrow (cohort 2) into 5 to 6-week-old female NBSGW mice. BM human cell engraftment was monitored by flow cytometry at 16 weeks post-transplant.

**Competitive bone marrow transplant experiments**

Primary transplant was performed after 4 weeks of control or alcohol liquid diet. 2.5 million BM cells from the donor groups (the young control (YC), young alcohol (YA), old control (OC), or old alcohol (OA)) were mixed with 2.5 million CD45.1 competitor cells in 100 μL PBS and retroorbitally injected into lethally irradiated (1000 cGy) 8-week-old female CD45.2 recipients. Secondary transplant was performed by retro-orbital injection of 5 million BM cells from primary transplant recipients into lethally irradiated secondary recipients. Engraftment was monitored every 4 weeks until 4-5 months for the primary transplant and until 10-12 weeks for the secondary transplant.

**LPS challenge and EdU incorporation assay**

LPS challenge was done either at 4-6 weeks post primary transplant or 12 weeks post secondary transplantation by injecting 5 mg/kg LPS O55:B5 from *E. coli* and Ethynyl-deoxy-uridine (EdU, Chemcruz) at 24 and 18 hours before sacrifice, respectively. EdU was detected using ClickiT Plus EdU Flow Cytometry kit following manufacturer's instructions (Thermofisher).

**Cell harvest and flow-cytometry analysis**

Peripheral blood (PB) was obtained from retroorbital bleeding and incubated with 2% dextran at 37 $^{o}$C. Spleen was ground on a 40 μm cell strainer and washed with fluorescence-activated cell sorting (FACS) buffer. Bone marrow was collected by cutting the end part of the tibiae, femurs, pelvis, and humeri, followed by centrifugation at 10,000 g for 20 seconds. Alveolar macrophages were collected via bronchoalveolar lavage as described previousy[6,7]. Erythrocytes were lysed with 1 mL ACK lysis solution for 15-20 minutes at room temperature. Cells were counted and stained either with Live Dead Fixable Yellow or Near-Infrared (NIR) dye as indicated. To analyze human xenotransplant cells in NBSGW mice, we used antibodies against hCD45, mCD45, CD33, CD19, CD24, CD14, CD15, CD41a diluted in FACS buffer for PB. To identify human HSPCs in NGSGW mice, we used antibodies against markers such as hCD45, mCD45, CD34, CD38, CD90, CD45RA, CD10, and CD135, diluted in FACS buffer.

For analysis of murine immunocytes in the spleen, BM, and PB, cells were stained with antibodies against Ter119, CD3e, B220, NK1.1, CD4, CD8a, CD11b, Ly6G, Ly6C, CD172a, CD11c, Siglec-F, and XCR1. For the analysis of the murine BM HSPC compartment, we stained cells with Lineage cocktails (CD11b, CD3e-, CD4, CD8a, Gr1, Ly6C, Ter119, B220), CD34, CD16/32, CD150, CD48, CD135, CD41, CD117, and Sca-1. When required, we added antibodies for CD61, CD63, MHC II, CD74, CD45.1, and CD45.2 (Table S5). For all specific information regarding antibody dilutions, clones, and fluorescence tags, please refer to Table S9. Cells were incubated with antibodies for 60 minutes on ice. Cell cycle activity and DNA damage were stained with anti-mouse Ki67 and anti-H2AX following fixation and permeabilization after cell surface staining. Data were acquired with an Aurora Cytek and analyzed with FlowJo v10 (TreeStar).

**Isolation and culture of human BM CD34+ cells**

Mononuclear cells were isolated by Ficoll gradient isolation from deidentified healthy donor BM cells. Human CD34+ cells were selected using either the SepMate protocol (StemCell Technologies) or a magnetic bead-based column (Miltenyi). Cells were viably frozen down or cultured in StemSpan SFEM II (StemCell Technologies, 09655) medium supplemented with 100 ng/ml of recombinant human FLT3 ligand (rhFLT3, Peprotech, 300-19), thrombopoietin (rhTPO, Peprotech, 300-18), and stem cell factor (rhSCF, Peprotech, 300-07), and 1x penicillin-streptomycin-glutamine (Gibco, 1078-016).

**Culture of human cell lines**

Human leukemia cell lines MOLM13 (*Deutsche Sammlung von Mikroorganismen und Zellkulturen GmbH*, DSMZ), were cultured in RPMI1640 medium supplemented with 10% FBS, 1% Penicillin/Streptomycin. Cells were seeded at a density of 500,000 cells/ml and split every 2-3 days. For adherent HeLa cell line (American Type Culture Collection, ATCC), cells were cultured in DMEM high glucose added with the same supplement and passaged every 2-3 days until reaching 80-90% confluency.

**dsRNA immunofluorescence staining**

Harvested murine alveolar macrophages, human leukemia cell lines, and human $CD34^+$ BM cells were harvested from *in vitro* culture, cytospun, and fixed with 4% paraformaldehyde on defrosted glass slides. Cells were then washed with PBS 3 times, permeabilized with 0.5% Triton-X 100 diluted in PBS and blocked using 5% FBS in PBS for 1 hour at room temperature. Cells were then stained with mouse anti-dsRNA J2 antibody (1:1000, Scicons, 10010200) at 4°C overnight in a humidified chamber. Following incubation, cells were washed for 5 minutes with blocking buffer 3 times to remove unbound primary antibody. Secondary antibody staining was performed using goat anti-mouse Alexa-594-conjugated IgG (1:2000 dilution, Thermo Scientific) for 1 hour at room temperature. Cells were imaged using EVOS M5000, and the immunofluorescence image was processed in FIJI/ImageJ using own-developed code, followed by quantification of dsRNA foci in CellProfiler using own-developed code (see code availability section).

**scRNA-seq of human $CD34^+$ $CD45^+$ and murine LSK cells**

Following cell sorting of human CD45+CD34+ bone marrow cells, the cells were processed for scRNA-seq using Chromium Next GEM Single Cell 3' Reagent Kits v3.1 (10X Genomics) according to the manufacturer's protocol, and libraries were then sequenced using Novaseq 6000 (Illumina) at the Single and Transcriptomics Core Facility of the Johns Hopkins School of Medicine. For each group of mice, 30,000-50,000 sorted murine live LSK cells were resuspended in FACS buffer (PBS, 0.5% FBS, 2 mM EDTA) and subsequently processed using Chromium Next GEM Single Cell 3' Reagent Kits v3.1 (10X Genomics) according to the manufacturer's protocol. Libraries were then sequenced using Novaseq 6000 (Illumina). The procedures were performed by the Sidney Kimmel Comprehensive Cancer Center's Experimental and Computational Genomics Core.

**scATAC-seq of murine LSK cells**

For each group, 100.000 LSK cells were centrifuged at cold temperature (300 g x 5 minutes). The supernatant was removed carefully, leaving only a tiny, translucent cell pellet. Pellet was resuspended in 50 μL PBS + 0.04% BSA and proceeded to the low-input nuclei isolation protocol for scATAC-seq according to 10X Genomics' instructions. At the final step, nuclei were resuspended in 10 μL nuclei buffer, resulting in more than 4,000 nuclei/μL to target 10,000 nuclei to be successfully sequenced. The scATAC-seq library was prepared using the Chromium Next GEM Single Cell ATAC Reagent Kits v2 (10X Genomics) according to the manufacturer's instructions. The resulting libraries were sequenced using NovaSeq 6000 (Illumina). All the scATAC-seq procedures were performed in the Single and Transcriptomics Core Facility of the Johns Hopkins School of Medicine.

**dsRNA immunoprecipitation (dsRIP-seq)**

For dsRIP-seq of human cell lines, we performed dsRNA immunoprecipitation according to the published protocol from the Tebaldi lab[8]. Forty million MOLM13 or HeLa cells were resuspended in 600 μL dsRIP buffer (see buffer list) supplemented with protease and RNase inhibitors, vortexed vigorously, and incubated for 5 minutes on ice. The lysate was equally divided into either J2 or IgG tubes, containing either 5 µg of J2 (Scicons, 10010200) or normal mouse IgG (Sigma, 12-371), respectively. They were incubated for 2 hours with rotation in the cold room. The J2-bound or non-specific IgG-bound RNAs were captured by incubating lysates with Dynabeads Protein G for 1 hour at 4 $^{o}$C. Beads were washed and the dsRNAs were extracted using TRIzol LS according to the manufacturer's instructions.

For dsRNA IP from mouse bone marrow cells, we followed the protocol used in this publication with some modifications[9]. Briefly, upon harvesting bone marrow cells from old alcohol or old control mice, we immunoprecipitated dsRNA from total RNA by using J2 antibody. We added 10 µg of J2 antibody to 50 µg of RNA diluted in dsRIP buffer, as mentioned above (see buffer list), and incubated overnight at 4 °C. The immunoprecipitated dsRNA was captured by incubating the lysates with Dynabeads Protein G for 4 hours at 4 °C. Beads were washed, and the dsRNAs were extracted from the beads using TRIzol (Invitrogen, 10296010) RNA extraction according to the manufacturer's protocol.

Library preparation was performed by using the KAPA RNA HyperPrep Kit with RiboErase HMR (Roche) according to the manufacturer's instructions. In brief, 50 ng of RNA were processed for rRNA depletion, which involved oligo hybridization with rRNA, digestion with RNase H, DNase digestion to remove the oligos, and cleanup with 2.2X SPRI beads. Following rRNA removal, the RNA was then converted into cDNA by performing 1$^{st}$ Strand Synthesis, 2$^{nd}$ Strand Synthesis, and A-tailing. At this step, the generated libraries were reverse-stranded. To reduce the impact of PCR amplification bias and as recommended by the Tebaldi lab for TE analysis[8], libraries were ligated with KAPA Universal UMI adapter (Roche) at 1.5 μM, diluted from a 33 μM stock in 10 mM Tris-HCl, pH 8.0. Ligated libraries were then purified using 0.7X SPRI beads and proceeded to library amplification by using KAPA UDI Primer mixes that contain sequencing barcode combinations. Libraries were quantified using the NEBNext Library Quant Kit for Illumina (NEB, E7630L). Libraries were adjusted to 4 nM equimolar for each sample, pooled together, and sequenced using Novaseq X (Illumina) at the Genetic Resources Core Facility (RRID: SCR_018669).

**Bioinformatics Analysis**

**Human xenotransplanted CD34+ scRNA-seq**

scRNA-seq analysis of xenotransplanted human CD34+ cells was performed using Seurat's v. 5.10. After filtering out a cluster of cells with low UMI counts and high mitochondrial gene expression, a total of 61,362 cells were processed with standard Seurat workflow and integrated through harmony by batch correction. PCA was performed to compute 50 principal components (PCs), clusters were found using the Louvain algorithm, and UMAP was plotted based on 50 PCs. The annotation was done with reference to HCA Bone Marrow Viewer[10], Azimuth v0.5.0 and CellTypist (Immune_All_Low)[11-13]. Differentially expressed genes (DEGs) were found using the FindMarkers function in Seurat, which computes significance through the Wilcoxon Rank

Sum test. DEGs were also identified after pseudobulking cells by group and cell type and subsequently compared between the alcohol and control group using DESeq2 v1.44.0 with default parameters. GSEA was performed using MSigDB's hallmark gene sets. All the computation was done in R. v4.4.1

**Murine LSK scRNA-seq**

CellRanger v.80 (10X Genomics) was used to demultiplex, filter, and align reads to mm39 mouse reference genome. BAM files generated by CellRanger were subsequently used for TE and gene counting using scTE package[14]. To this aim, we first generated a scTE index file by running the scTE_build function using the mm39 repeat masker bed file (UCSC) and the Genecode gtf annotation file (Gencode). Then, we called cell barcodes (CB), and used these barcodes to filter reads based on cell barcodes (see code availability). The filtered BAM files were run for scTE to generate h5ad countmatrix file containing genes and TEs.

The resulting h5ad files were loaded and converted into Seurat objects using Seurat v.5.10[15]. All the Seurat objects were merged, and high-quality reads were filtered based on UMI count (nUMI) ≥ 500, library complexity (nGene) ≥ 250, and library complexity (log10GenesPerUMI) ≥ 0.80. Since alcohol can also affect mitochondria, we did not perform direct filtering based on mitochondrial reads but rather based on the cutoffs, which will indirectly remove low-quality cells with high mitochondrial transcripts. The filtered libraries were then transformed and normalized using SCTransform v0.4.1, utilizing variance-stabilized transformation (vst) v2. The integration of several libraries was performed using the CCA integration algorithm. Following integration, the principal component analysis (PCA) value was determined using ElbowPlot. We used 20 PCA to perform clustering using the Louvain algorithm in the FindCluster() function, and with cluster resolution 0.4. Uniform Manifold Approximation clusters were visualized by running the RunUMAP() function using the same PCA value. Cluster marker genes were identified using the FindMarker() function. Finally, the HSPC clusters were identified manually by cross-checking with published HSPC markers from the following publications. Differential gene expression analysis was done in bona-fide LT-HSC cluster using FindMarkers() function by comparing two groups using Wilcoxon-ranked sum test; YA vs. YC, OA vs. OC, and OC vs. YC, and the significantly differentiated genes were filtered based on padj ≤ 0.05 and absolute avg_log2FC > 0.58 and plotted using EnhancedVolcano package. To identify enriched pathways, we used fgsea package using hallmarks, reactome, and gene ontology biological processes gene sets obtained from molecular signature databases (MSigDb) [16]. To do this, we first filtered genes with pVal < < 0.001 and ranked them by avg_log2FC, which was then input into the fgsea() function. The top enriched pathways were visualized in a dotplot. Module scores were calculated using Seurat's AddModuleScore() function by inputting previously published module genes. All the Seurat analyses were performed in R version 4.2.1 (https://www.r-project.org) on the Linux Green Obsidian computation workstation.

**Murine LSK scATAC-seq**

The generated sequencing reads were preprocessed and aligned using CellRanger-ATAC v.2.1.0 (10X Genomics). Reads were aligned into the mm10 reference genome from 10X Genomics's cellranger-arc-mm10-2020-A-2.0.0 to generate BAM files and count matrices. We used BAM files for differential open chromatin analysis using the ArchR v.1.0.3 package run in the R v.4.4.1 environment (https://www.r-project.org). We followed analytical procedures for ArchR as described in its original publication [17] and repository (https://www.archrproject.com/bookdown/index.html). Briefly, BAM files were loaded into ArchR to generate ArchR arrow and project files. QC metrics were plotted using the plotQC() function to show raw data enrichment for transcription start site (TSS) and number of fragments. Doublet enrichment scores were calculated using the addDoubletScores() function. We passed cells exhibiting doublet scores enrichment < 1.5, TSS enrichment > 6, and nFragment > 1000. We verified the batch correction with and without Harmony, which did not alter the clustering, and decided to proceed with the correction using Harmony. Using iterative latent semantic indexing (LSI) and harmony correction, we identified 22 clusters, which were then refined into 14 clusters via cluster label transfer from scRNA-seq clusters and manual cross-checking of marker gene accessibility. We converted the mm10 RepeatMasker bed file into genomic ranges to dissect TE-overlapping genomic regions, as previously described[18]. Peak was called using MACS2 v.2.2.9.1 peak caller; the resulting peak list was added to the ArchR project by running the addPeakMarker() function. Differential peak analysis to identify cluster-specific peaks within global UMAP or within LT-HSC subsets comparing OA vs. OC was done based on FDR < 0.05 and Log2FC > 0.58. Transcription factor (TF) motif analysis was done to identify cluster-specific TF enrichment, as well as within LT-HSC subsets comparing OA vs. OC. Over-representation analysis (ORA) was done using rGREAT v.2.5.4 software[19] (https://jokergoo.github.io/rGREAT/index.html) run locally in R environment using gene ontology biological processes obtained from MSigDB database. To reduce redundancy in GO enrichment results, semantically similar GO terms were clustered using rrvgo[20], and one representative term per cluster was selected based on fold enrichment. All reported statistics (fold enrichment, FDR, region hits) were derived from the original GREAT results.

**Murine bone marrow dsRIP-seq**

FASTQ files were preprocessed using a computational pipeline based on the KAPA Universal UMI Adapter, with modifications tailored for RNA-seq. In brief, the FASTQ files were merged and converted into unmapped BAM files using Picard. Then the UMIs were extracted from unmapped BAM files using Fgbio tools to generate UMI-extracted unmapped BAM files, which were then converted back into two FASTQ files per library by using Picard. Then, the FASTQ files were filtered and trimmed using the fastp program. The trimmed FASTQ files were then aligned into the mm10 reference genome using STAR v2.7.10a. Following alignment, the BAM files were then deduplicated from UMIs by using UMI-tools[21]. Reads counting was done using the featureCount function from the Subread package[22] to generate gene and TE count matrices using gene and TE annotation files downloaded from Molly Hammel's lab (https://hammelllab.labsites.cshl.edu/software/)[23]. Differential expression analysis was performed using DESeq2[24] to identify differentially enriched genes and TEs in J2-RIP samples. We also performed calculations based on the published pipeline from the Tebaldi lab to identify enrichment of dsRIP genes, using the input as a reference[8]. The analysis was done in R v4.2.1 (https://www.r-project.org)

#### Supplemental Table Legends

**Table S1**. DEG List in murine LT-HSCs

**Table S2**. GSEA Pathway List in murine LT-HSCs

**Table S3** DE TE list in murine LT-HSC

**Table S4**. dsRNA gene lists murine BM

**Table S5**. Gene Ontology Total OCRs

**Table S6**. Gene_Ontology TE-overlapping OCRs

**Table S7**. TF motif list for UpSet Plot

**Table S8**. TF motifs enriched in Total and TE OCRs

**Table S9.** Flow-cytometry panels

**Table S10.** Key Resources Table

**Table S9. Flow-cytometry panels**

Murine immune panel for PB, spleen and BM samples

| No | Marker | Conjugate | Clone | Vendor | Catalog | Dilution | RRID |
|---|---|---|---|---|---|---|---|
| 1 | CD3e | PE | 145-2C11 | BD | 553064 | 1 in 100 | AB_394597 |
| 2 | B220 | FITC | RA3-6B2 | BD | 553088 | 1 in 100 | AB_394618 |
| 3 | Ter119 | APC | TER-119 | eBioscience | 17-5921-81 | 1 in 1200 | AB_469472 |
| 4 | CD11b | BUV395 | M1/70 | BD | 565976 | 1 in 100 | AB_2721166 |
| 5 | Ly6C | BV480 | HK1.4 | BD | 569438 | 1 in 200 | AB_3685063 |
| 6 | Ly6G | eF450 | 1A8 | BD | 562737 | 1 in 200 | AB_2737756 |
| 7 | CD4 | R718 | GK1.5 | BD | 567311 | 1 in 200 | AB_2916549 |
| 8 | CD8 | RB780 | 53-6.7 | BD | 568692 | 1 in 100 | AB_2936840 |
| 9 | NK1.1 | PerCP-cy5.5 | PK136 | BD | 551114 | 1 in 200 | AB_394052 |
| 10 | CD172a | BUV496 | P84 | BD | 741131 | 1 in 200 | AB_2870711 |
| 11 | CD11c | PE-CF594 | N418 | BD | **565591** | 1 in 200 | AB_2869692 |
| 12 | MHCII | BUV737 | M5/114.15.2 | BD | 569176 | 1 in 200 | AB_3684842 |
| 13 | Siglec-F | APC/Cy7 | E50-2440 | BD | 565527 | 1 in 200 | AB_2732831 |
| 14 | CD45 | BV786 | 30-F11 | BD | 564225 | 1 in 200 | AB_2716861 |
| 15 | XCR1 | BV650 | ZET | Biolegend | 148220 | 1 in 200 | AB_2566410 |
| 16 | F480 | PE-Cy7 | BM8 | Biolegend | 123114 | 1 in 200 | AB_893478 |

LSK sorting panel for murine BM

| No | Marker | Tag | Clone | Vendor | Catalog | Dilution | RRID |
|---|---|---|---|---|---|---|---|
| 1 | cKit | APC | 2B8 | ThermoFisher | 17-1171-82 | 1 in 100 | AB_469430 |
| 2 | Sca-1 | BV421 | D7 | Biolegend | 115904 | 1 in 100 | AB_313683 |
| 3 | CD3e | PE | 145-2C11 | BD Pharmingen | 563565 | 1in 100 | AB_2738278 |
| 4 | CD4 | PE | Rm4-5 | BD Pharmingen | 740208 | 1 in 100 | AB_2734761 |
| 5 | CD8a | PE | 53-6.7 | BD Pharmingen | 563786 | 1 in 100 | AB_2732919 |
| 6 | B220 | PE | RA3-6B2 | BD Pharmingen | 563793 | 1 in 100 | AB_2738427 |
| 7 | CD11b | PE | M1/70 | BD Pharmingen | 563553 | 1 in 100 | AB_2738276 |
| 8 | Gr1 | PE | RB6-8C5 | BD Pharmingen | 553128 | 1 in 100 | AB_394644 |
| 9 | Ter119 | PE | Ter-119 | BD Pharmingen | 563827 | 1 in 100 | AB_2738438 |

Murine HSPC surface marke panel

| No | Marker | Tag | Clone | Vendor | Catalog | Dilution | RRID |
|---|---|---|---|---|---|---|---|
| 1 | CD34 | FITC | RAM34 | ThermoFisher | 11-0341-82 | 1 in 100 | AB_465021 |
| 2 | cKit | APC | 2B8 | ThermoFisher | 17-1171-82 | 1 in 100 | AB_469430 |
| 3 | CD16/CD32 | AF700 | 93 | ThermoFisher | 56-0161-82 | 1 in 100 | AB_493994 |
| 4 | CD48 | BV510 | HM48-1 | BD Pharminger | 563536 | 1 in 100 | AB_2738266 |
| 5 | CD150 | PE | TC15-12F12.1 | Biolegend | 115904 | 1 in 100 | AB_313683 |
| 6 | Sca-1 | BV650 | D7 | Biolegend | 108143 | 1 in 100 | AB_2629684 |
| 7 | CD135 (Flk2) | BV421 | A2F10.1 | BD Bioscience | 562898 | 1 in 50 | AB_2737876 |
| 8 | MHCII | BUV737 | M5/114.15.2 | BD | 569176 | 1 in 200 | AB_3684842 |
| 9 | CD63 | PE-Cy7 | NVG2 | Biolegend | 143909 | 1 in 200 | AB_2565499 |
| 10 | CD61 | PE-Dazzle | 2C9-G2 | Biolegend | 104321 | 1 in 200 | AB_2813932 |
| 11 | CD74 | BV711 | In-1 | BD Bioscience | 743735 | 1 in 100 | AB_2741708 |
| 12 | CD41 | RB780 | MWReg30 | BD Pharmingen | 755714 | 1 in 100 | AB_3688041 |
| 13 | CD3e (Lin) | BUV395 | 145-2C11 | BD Pharmingen | 563565 | 1in 100 | AB_2738278 |
| 14 | CD4 (Lin) | BUV395 | Rm4-5 | BD Pharmingen | 740208 | 1 in 100 | AB_2734761 |
| 15 | CD8a (Lin) | BUV395 | 53-6.7 | BD Pharmingen | 563786 | 1 in 100 | AB_2732919 |
| 16 | B220 (Lin) | BUV395 | RA3-6B2 | BD Pharmingen | 563793 | 1 in 100 | AB_2738427 |
| 17 | CD11b (Lin) | BUV395 | M1/70 | BD Pharmingen | 563553 | 1in 100 | AB_2738276 |
| 18 | Gr1 (Lin) | BUV395 | RB6-8C5 | BD Pharmingen | 566218 | 1 in 100 | AB_2739608 |
| 19 | Ter119 (Lin) | BUV395 | Ter-119 | BD Pharmingen | 563827 | 1 in 100 | AB_2738438 |

Murine HSPC panel with Ki67 and γH2AX

| No | Marker | Tag | Clone | Vendor | Catalog | Dilution | RRID |
|---|---|---|---|---|---|---|---|
| 1 | CD34 | FITC | RAM34 | ThermoFisher | 11-0341-82 | 1 in 100 | AB_465021 |
| 2 | cKit | APC | 2B8 | ThermoFisher | 17-1171-82 | 1 in 100 | AB_469430 |
| 3 | CD16/CD32 | AF700 | 93 | ThermoFisher | 56-0161-82 | 1 in 100 | AB_493994 |
| 4 | CD48 | BV510 | HM48-1 | BD Pharminger | 563536 | 1 in 100 | AB_2738266 |
| 5 | CD150 | PE | TC15-12F12.1 | Biolegend | 115904 | 1 in 100 | AB_313683 |
| 6 | Sca-1 | BV650 | D7 | Biolegend | 108143 | 1 in 100 | AB_2629684 |
| 7 | CD135 (Flk2) | BV421 | A2F10.1 | BD Bioscience | 562898 | 1 in 50 | AB_2737876 |
| 8 | CD3e (Lin) | BUV395 | 145-2C11 | BD Pharmingen | 563565 | 1in 100 | AB_2738278 |
| 9 | CD4 (Lin) | BUV395 | Rm4-5 | BD Pharmingen | 740208 | 1 in 100 | AB_2734761 |
| 10 | CD8a (Lin) | BUV395 | 53-6.7 | BD Pharmingen | 563786 | 1 in 100 | AB_2732919 |
| 11 | B220 (Lin) | BUV395 | RA3-6B2 | BD Pharmingen | 563793 | 1 in 100 | AB_2738427 |
| 12 | CD11b (Lin) | BUV395 | M1/70 | BD Pharmingen | 563553 | 1in 100 | AB_2738276 |
| 13 | Gr1 (Lin) | BUV395 | RB6-8C5 | BD Pharmingen | 566218 | 1 in 100 | AB_2739608 |
| 14 | Ter119 (Lin) | BUV395 | Ter-119 | BD Pharmingen | 563827 | 1 in 100 | AB_2738438 |
| 15 | Ki67 | BV711 | B56 | BD Bioscience | 563755 | 1 in 50 | AB_2738406 |
| 16 | γH2AX | PE-CF594 | N1-431 | BD Bioscience | 564719 | 1 in 50 | AB_2738913 |

Peripheral blood panel for human xenotransplant

| No | Marker | Tag | Clone | Company | Catalog | Dilution | RRID |
|---|---|---|---|---|---|---|---|
| 1 | hCD45 | BV510 | HI30 | BD Biosciences | 563204 | 1 in 200 | AB_2738067 |
| 2 | mCD45 | BV650 | 30-F11 | BD Biosciences | 563410 | 1 in 100 | AB_2738189 |
| 3 | hCD33 | APC | WM53 | BD Biosciences | 551378 | 1 in 200 | AB_398502 |
| 4 | hCD19 | PE-CY5 | HIB19 | BD Biosciences | 555414 | 1 in 100 | AB_395814 |
| 5 | hCD24 | PE | ML5 | BD Biosciences | 555428 | 1 in 100 | AB_395822 |
| 6 | hCD14 | AF700 | M5E2 | BD Biosciences | 557923 | 1 in 100 | AB_396944 |
| 7 | hCD15 | PE-Cy7 | HI98 | BD Biosciences | 560827 | 1 in 200 | AB_10563901 |
| 8 | hCD41a | V450 | HIP8 | BD Biosciences | 561425 | 1 in 500 | AB_10714784 |

HSPC panel for human xenotransplant

| No | Marker | Tag | Clone | Company | Catalog | Dilution | RRID |
|---|---|---|---|---|---|---|---|
| 1 | hCD45 | BV510 | HI30 | BD Biosciences | 563204 | 1 in 100 | AB_2738067 |
| 2 | mCD45 | BV650 | 30-F11 | BD Biosciences | 563410 | 1 in 100 | AB_2738189 |
| 3 | CD34 | APC/Fire750 | 581 | Biolegend | 343536 | 1 in 50 | AB_2650736 |
| 4 | CD38 | APC | HB7 | BD Biosciences | 567144 | 1 in 500 | AB_2916467 |
| 5 | CD90 | BV421 | 5E100 | BD Biosciences | 562556 | 1 in 200 | AB_2737651 |
| 6 | CD45RA | PE | HI100 | BD Biosciences | 555489 | 1 in 500 | AB_395880 |
| 7 | CD10 | PE-Cy7 | HI10a | BD Biosciences | 565282 | 1 in 200 | AB_2739153 |
| 8 | CD135 | BV711 | 4G8 | BD Biosciences | 563908 | 1 in 100 | AB_2738479 |

**Table S10. Key Resources Table**

Immunohistochemistry and dsRIP-seq antibodies

| No | Antibody | Host | Company | Catalog | RRID |
|---|---|---|---|---|---|
| 1 | J2 | Mouse | Scicons | 10010200 | AB_2651015 |
| 2 | Normal mouse IgG | Mouse | MilliporeSigma | 12-371 | AB_145840 |

Critical assay kits

| No | Kits | Vendor | Catalog |
|---|---|---|---|
| 1 | EasySep Human CD34 Positive Isolation Kit II | StemCell Technologies | #17856 |
| 2 | Qubit RNA HS Assay Kit | Invitrogen | Q32855 |
| 3 | KAPA RNA HyperPrep Kit | Roche | #0810593 |
| 4 | KAPA RIboErase Kit | Roche | #07962266001 |
| 5 | KAPA Universal UMI Adapter | Roche | #09329862001 |
| 6 | NEBNext Library Quantification Kit | New England Biolab | E7630L |
| 7 | FAST SYBR Green Master Mix | Applied Biosystems | #4385612 |
| 8 | SuperScript™ III First-Strand Synthesis System | Invitrogen | #18080051 |
| 9 | ClickiT EdU Alexa Fluor 647 Flow Cytometry | Invitrogen | C10424 |

Human cell lines

| No | Cell lines | Vendor | Catalog |
|---|---|---|---|
| 1 | HeLa | ATCC | CCL-2 |
| 2 | MOLM-13 | DSMZ | ACC-554 |

Reagents and chemicals

| No | Reagents and chemicals | Vendor | Catalog |
|---|---|---|---|
| 1 | Recombinant human FLT3 ligand | Peprotech | AF-300-19 |
| 2 | Recombinant human thrombopoietin | Peprotech | 300-18-100UG |
| 3 | Recombinant stem cell Factor | Peprotech | 300-07 |
| 4 | Penicilin-streptomycin-glutamine | Gibco | 1078-016 |
| 5 | StemSpan SFEM II | StemCell Technologies | #09655 |
| 6 | RPMI + L-Glutamine | Gibco | 11875-093 |
| 7 | DMEM high glucose + GLUTAMAX | Gibco | 10569-010 |
| 8 | Sodium chloride 5M solution | Promega | V4221 |
| 9 | Magnesium chloride | Sigma-Aldrich | M1028-100ML |
| 10 | Tris-HCl, 1M solution, pH 7.4 | Quality Biologicals | 723949 |
| 11 | IGEPAL CA-630 | Sigma-Aldrich | 18896-50ML |
| 12 | cOmplete, Mini, EDTA-free Protease Inhibitor cocktail | Sigma-Aldrich | 11836170001 |
| 13 | SUPERase in RNase Inhibitor | Thermo Fisher Scientific | AM2696 |
| 14 | Dynabeads Protein G | Thermo Fisher Scientific | 10004D |
| 15 | TRIzol Reagent | Thermo Fisher Scientific | 15596026 |
| 16 | Chloroform molecular biology grade | MP Biomedicals | 194002 |
| 17 | GlycoBlue Coprecipitant | Thermo Fisher Scientific | AM9515 |
| 18 | Ethanol | Fisher Bioreagents | BP2818-500 |
| 19 | Lieber d'Carli Powder Mix Alcohol | Bio-Serv | F1258SP |
| 20 | Lieber d'Carli Powder Mix Control | Bio-Serv | F1259SP |
| 21 | ACK lysing solution | Quality Biologicals | 118-156-721 |
| 22 | D-PBS pH 7.2-7.4 | Gibco | 10010-023 |
| 23 | Molecular grade water | Corning | 46-000-CV |
| 24 | Cell Strainer 40 micron | Corning | 431750 |
| 25 | Fetal bovine serum | Biotechne | S11150 |
| 26 | BD Cytofix Fixation Buffer | BD Biosciences | 554655 |
| 27 | BD Horizon Brillian Stain Buffer | BD Biosciences | 566349 |
| 28 | Vectashield antifade mounting medium | VectorLab | H-1000 |
| 29 | 5% Digitonin | Invitrogen | BN2006 |
| 30 | Dextran solution from Leuconostoc 20% w/w | Sigma-Aldrich | D8802-50ML |
| 31 | NuPAGE 4-12% Bis-Tris Gel | Invitrogen | NP0335BOX |
| 32 | Halt Protease & Phosphatase Single-Use Inhibitor Cocktail 100X | Thermo Scientific | 78442 |
| 33 | BD Phosflow Perm Buffer III | BD Biosciences | 558050 |
| 34 | Lipopolysaccharides from E. coli O55:B5 | Sigma-Aldrich | L2637-5MG |

**Supplemental Figures**

**Figure S1. Chronic alcohol consumption slightly reduced the survival of female mice.**

**Figure S2. Chronic alcohol consumption alters the HSPC compartment by promoting replicative stress.**

**Figure S3. Identification of age-associated transcriptional changes in aged LT-HSCs.**

**Figure S4. Chronic alcohol consumption causes inflammaging phenotypes and increased p53 module scores in upstream progenitor subpopulations.**

**Figure S5. Chronic alcohol consumption upregulates TE in HSPCs**

**Figure S6. scATAC-seq cluster identification from LSK cells.**

**Figure S7. Chronic alcohol consumption increases chromatin accessibility of TEs.**

**Figure S8. Chronic alcohol consumption promotes myeloid bias in human CD34+ cells and induces the dsRNA-sensing pathway.**

#### Supplemental Figure 1

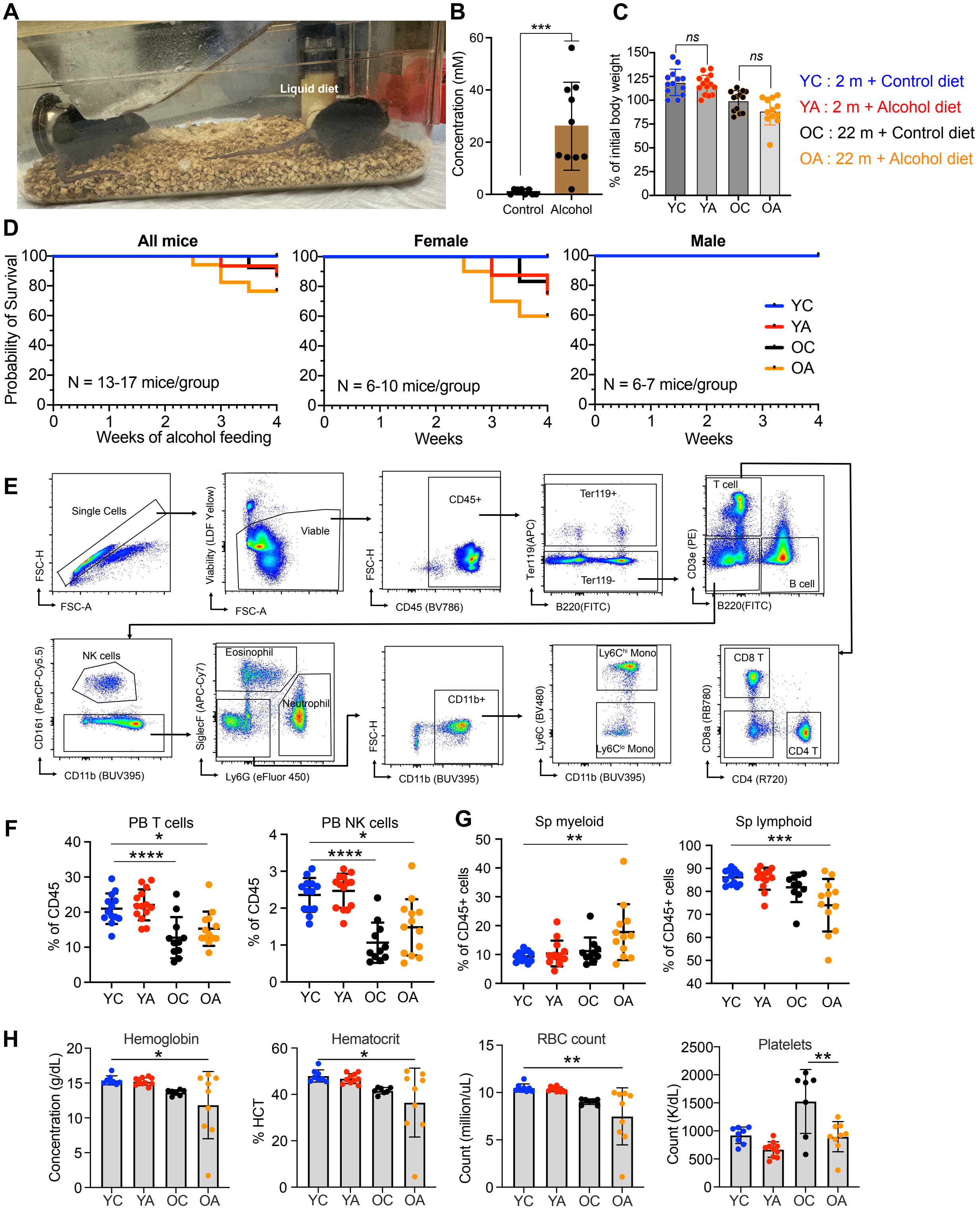


376

**Figure S1. Chronic alcohol consumption slightly reduced the survival of female mice.**

**(A)** A representative photo of an alcohol feeding cage.

**(B)** Blood ethanol concentration (BEC) measured in mM from YA and YC mice at the end of 4-5 weeks of alcohol feeding. Bars indicate Mean ± SD, dots represent individual mice, Student's t-test, *$p < 0.05$; **$p < 0.01$; ***$p < 0.0001$; ****$p < 0.0001$. Data were pooled from 2 independent experiments.

**(C)** Body weight changes as a percent of the initial body weight at the end of 4-5 weeks of alcohol feeding. Bars indicate mean ± SD. Dots represent individual mice, one-way ANOVA, *$p < 0.05$; **$p < 0.01$; ***$p < 0.0001$; ****$p < 0.0001$. Data were pooled from 3 independent experiments.

**(D)** Kaplan-Meier's survival curves of all mice and sex-stratified groups. Data were pooled from 3 independent experiments. YC, young control; YA, young alcohol; OC, old control; OA, old alcohol.

(**E**) Gating strategy to analyze immunocytes in the peripheral blood and spleen.

(**F**) PB CD45+ cell subpopulations.

(**G**) Quantification of myeloid and lymphoid cells within CD45+ spleen immunocytes.

(**H**) Complete blood count data showing hemoglobin concentration, percent hematocrit, RBC count, and platelets in mouse PB at 4 weeks of alcohol consumption.

**Supplemental Figure 2**

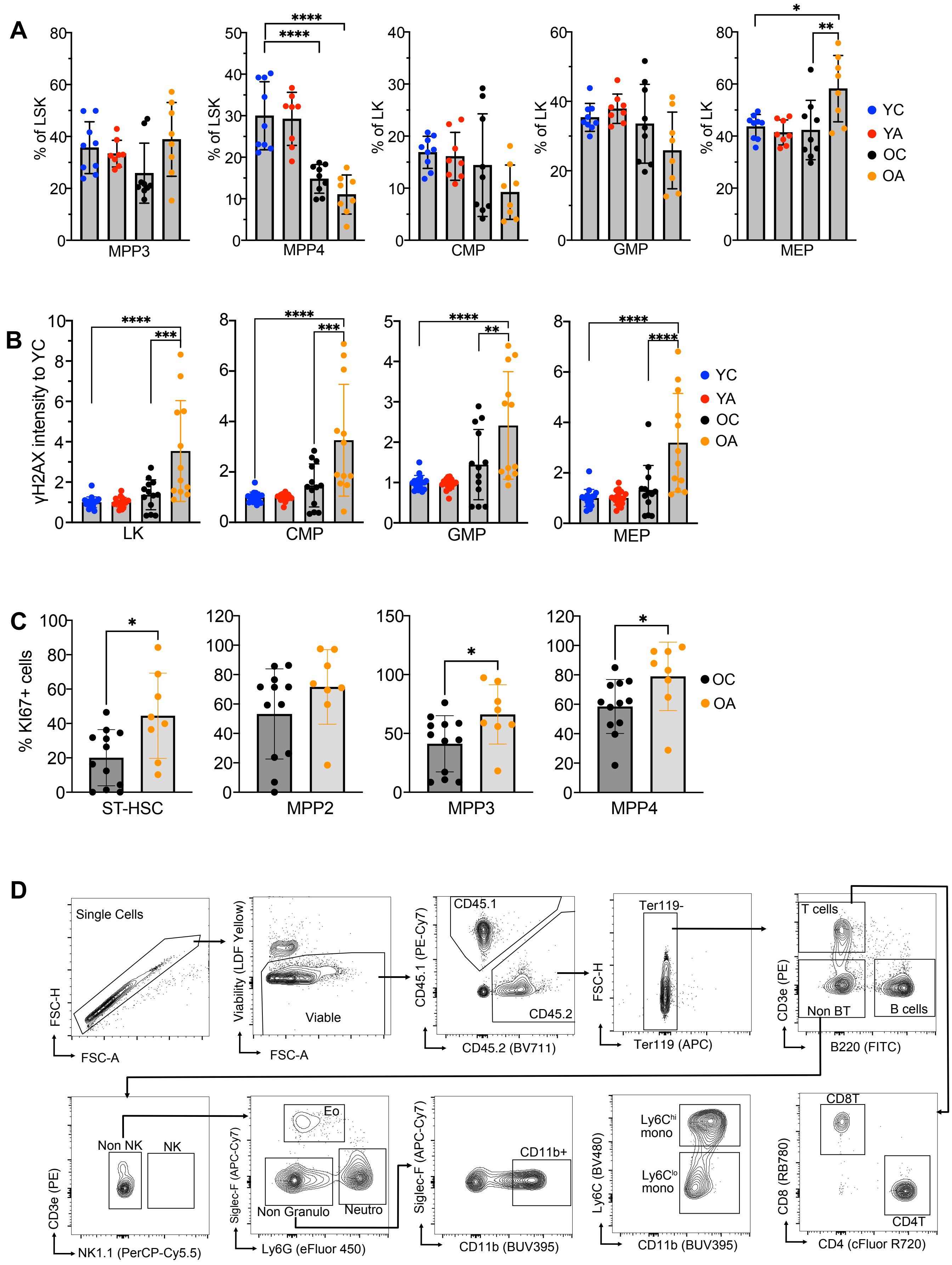

**Figure S2. Chronic alcohol consumption alters the HSPC compartment by promoting replicative stress.**

**(A)** Proportion of murine MPP3, MPP4, CMP, GMP, and MEP within parent LSK or LK stem progenitor cells, respectively, comparing 4 different groups.

**(B)** γH2AX intensity in LK progenitors comparing 4 different groups.

(**A-B**) One-way ANOVA followed by Tukey's multiple comparisons, *$p < 0.05$; **$p < 0.01$; ***$p < 0.001$; ****$p < 0.0001$. Dots represent individual mice. Bars represent mean ± SD. Data were pooled from 3 independent experiments.

**(C)** Percentage of Ki67+ cells within ST-HSCs and MPP2/3/4 at the 4-5 weeks alcohol feeding experiments. Student's t-test, *$p < 0.05$. Dots represent individual mice. Bars represent mean ± SD. Data were pooled from 3 independent experiments.

**(D)** Gating strategy to analyze donor-derived (CD45.2) immunocytes from a serial competitive bone marrow transplant experiment. MPP, multipotent progenitor; CMP, common myeloid progenitor; GMP, granulocyte-macrophage progenitor; MEP, megakaryocyte-erythroid progenitor; LSK, lineage-negative, Sca-1 positive, cKit positive; LK, lineage negative, cKit positive; ST-HSC, short-term hematopoietic stem cell.

#### Supplemental Figure 3

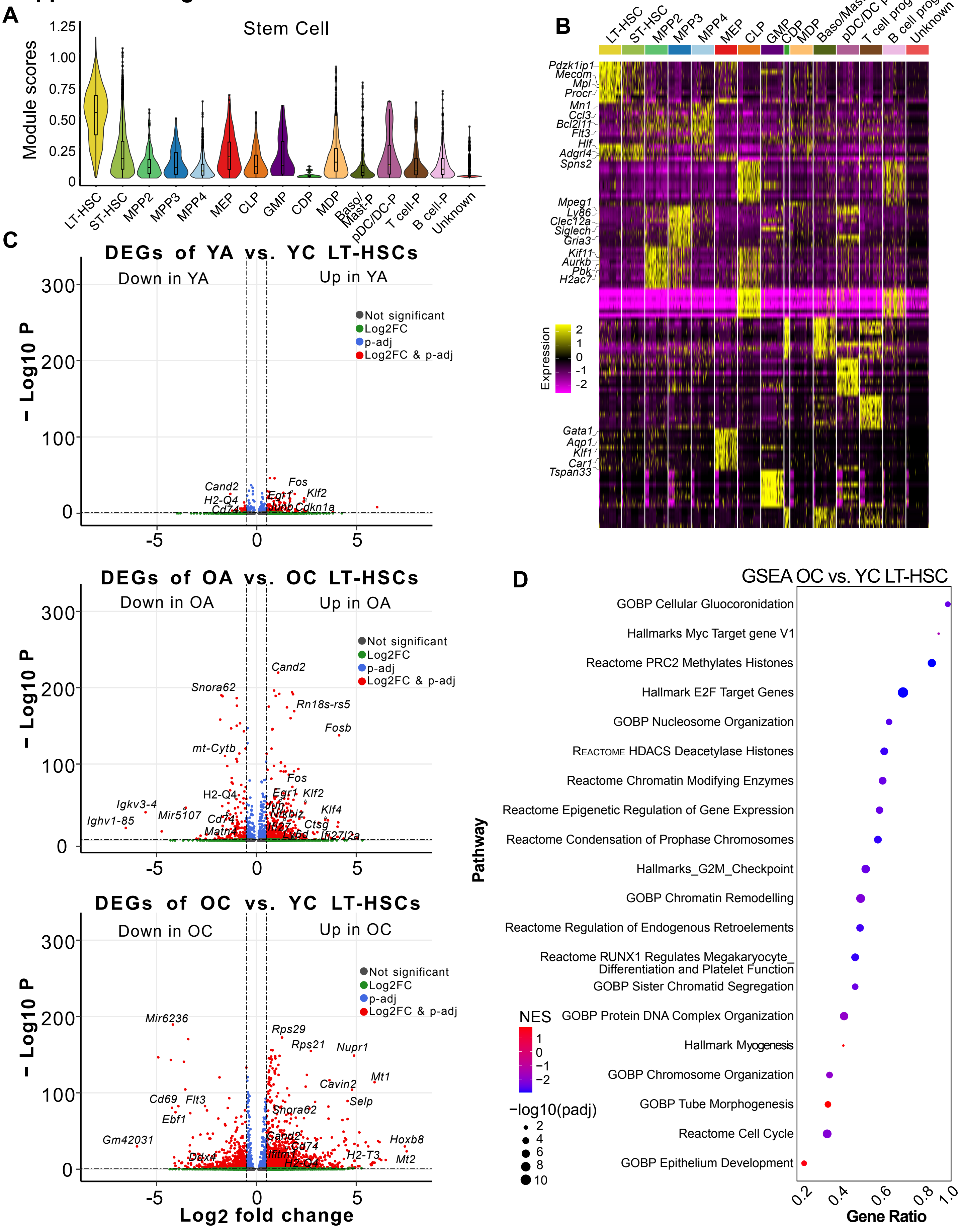

**Figure S3. Identification of age-associated transcriptional changes in aged LT-HSCs.**

**(A)** Stem cell module score calculated by using HSC genes reported by Nestorowa et al 2016[25].

**(B)** A heatmap showing the top 10 cluster marker genes calculated using the FindMarkers function and the Wilcoxon rank-sum statistical test with threshold *padj* < 0.01 and absolute log2FC > 1.6.

**(C)** Volcano plot depicting differentially expressed genes (DEGs) calculated using the Wilcoxon ranked-sum test. Red dots represent DEGs filtered based on false-discovery rate ≤ 0.05, log2FC ≥ 0.58. The full list of DEGs is provided in **Table S1.**

**(D)** Dot plot showing top pathways enriched in OC vs. YC LT-HSCs, with positive and negative normalized enrichment scores (NES) indicating upregulation and downregulation of the respective pathways. GSEA was calculated from differentially expressed genes that passed a p < 0.01 cutoff, which were subsequently ranked based on average Log2FC before calculation using the fgsea package. The full list of pathways is provided in **Table S2**

#### Supplemental Figure 4

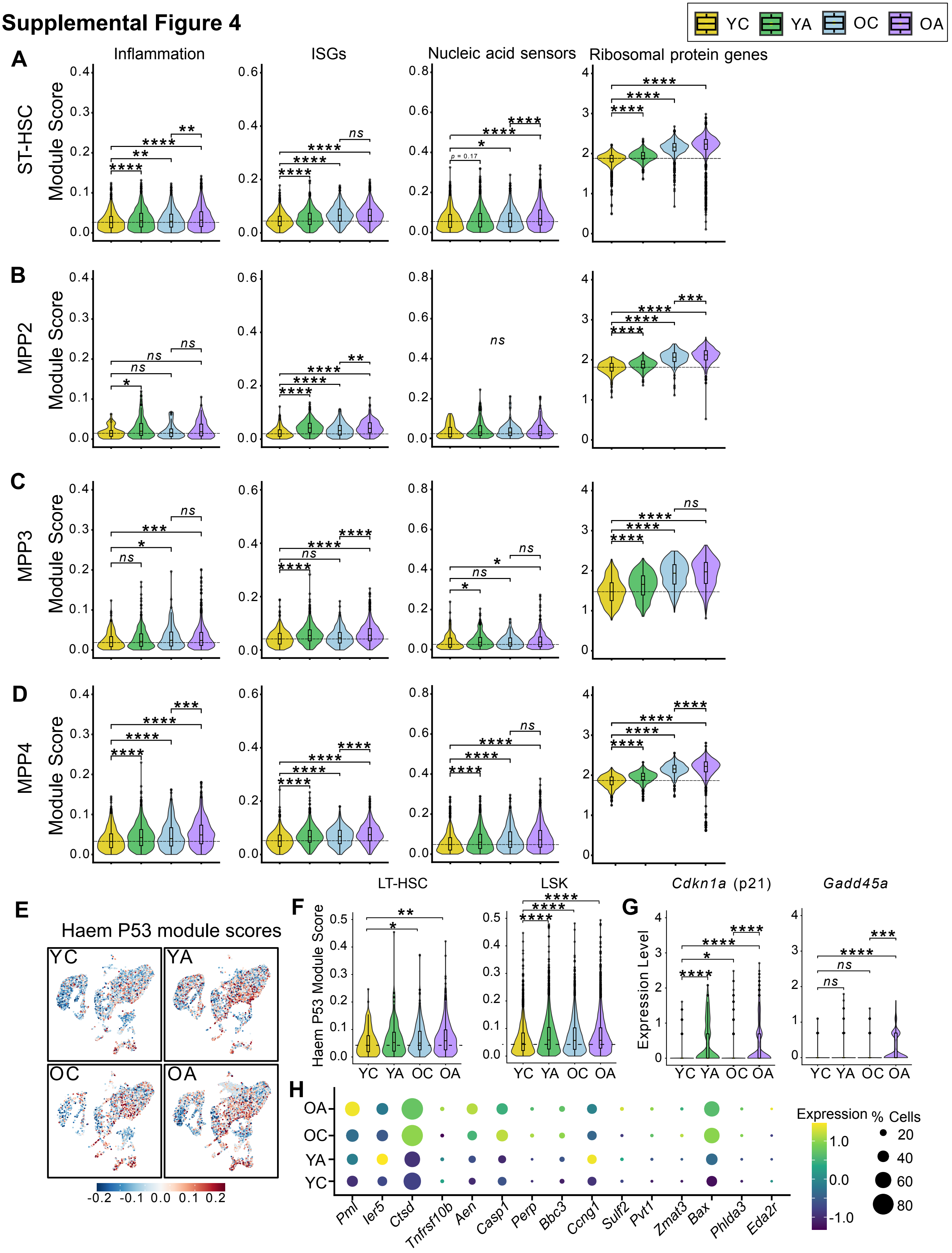

**Figure S4. Chronic alcohol consumption causes inflammaging phenotypes and increased p53 module scores in upstream progenitor subpopulations.**

(**A-D**) Violin plot showing module scores in inflammation, ISGs, nucleic acid sensors, and ribosomal protein genes in **(A)** ST-HSC, **(B)** MPP2, **(C)** MPP3, and **(D)** MPP4.

(**E**) Calculated Haem p53 module score projected onto UMAP, separated into 4 groups. Module score was calculated by using 16 p53 target genes known to be differentially expressed in HSPCs deficient in *Fancd2* and *Aldh2*[26].

**(F)** Violin plot showing Haem p53 module score in the LT-HSC cluster and all combined clusters in LSK.

**(G)** Violin plot showing the expression of *Cdkn1a* and *Gadd45a* genes across the 4 groups.

**(H)** Dot plot showing the expression levels of 15 Haem p53 genes in the LT-HSC cluster. Statistical analyses were performed using the Wilcoxon rank-sum test.

ISG, interferon-stimulated gene.

**Supplemental Figure 5**

**A**

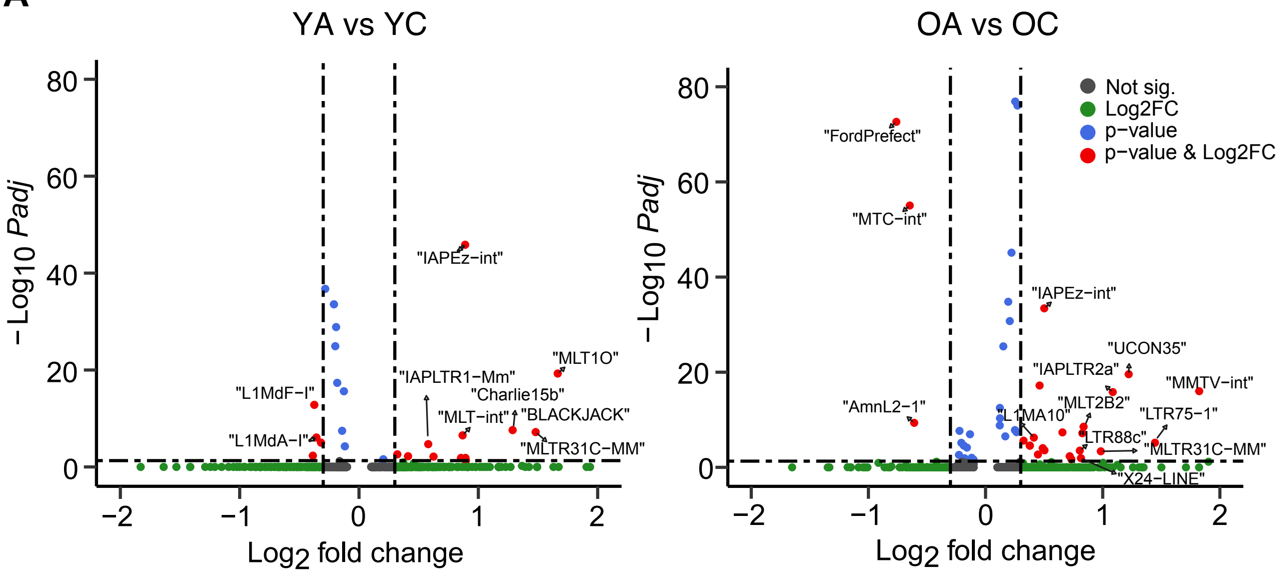


**B**

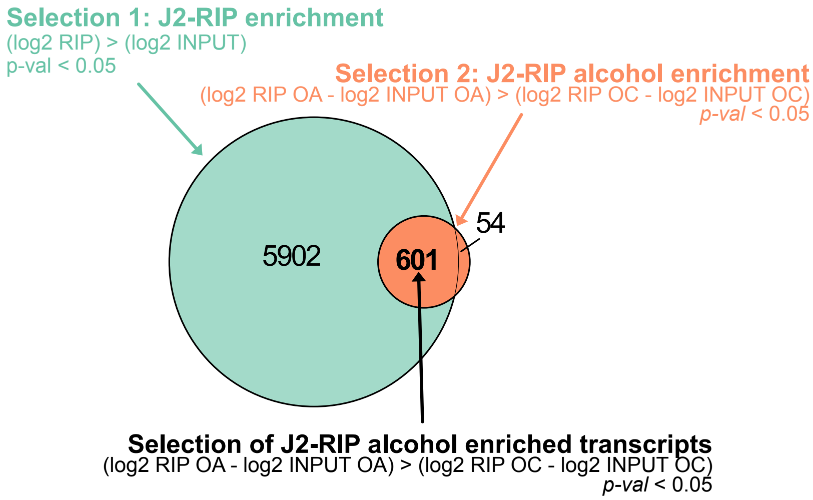


**C**

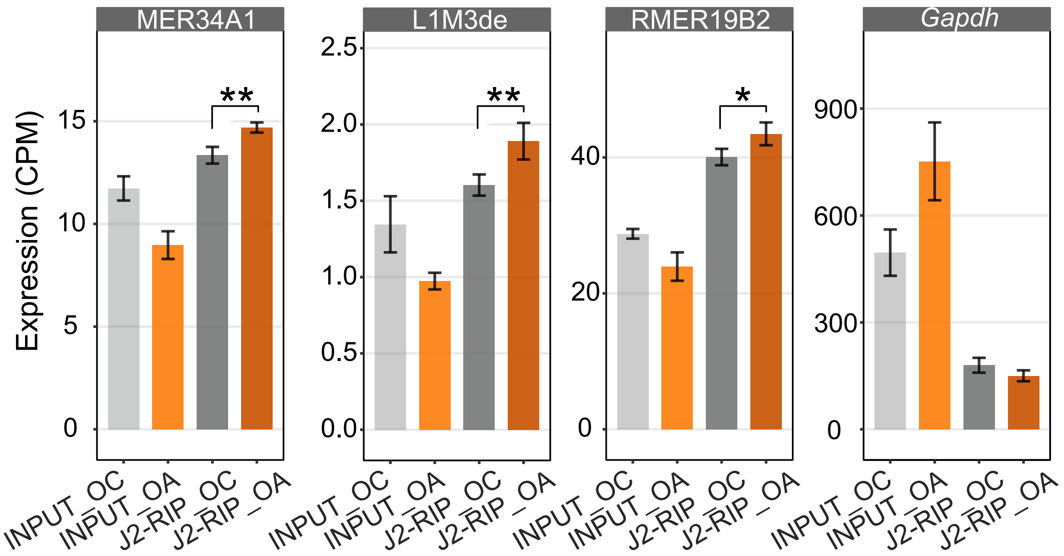

**Figure S5. Chronic alcohol consumption upregulates TE in HSPCs.**

(**A**) Volcano plots showing differentially expressed TEs (DE TEs) in LT-HSC based on YA vs. YC and OA vs. OC comparisons using padj ≤ 0.05 and [log2FC] ≥ 0.58. The full list of DE TEs is provided in **Table S3**

(**B**) Venn Diagram showing the number of dsRNA-enriched transcripts in OA vs. OC mice that passed selection criteria 1 and 2. The full list of dsRNA-enriched genes are provided in **Table S4**.

(**C**) Barplot showing count per million (cpm) expression of representative LTR and LINE species, as well as negative control *Gapdh* within dsRNAome (n = 5 mice/group).

#### Supplemental Figure 6

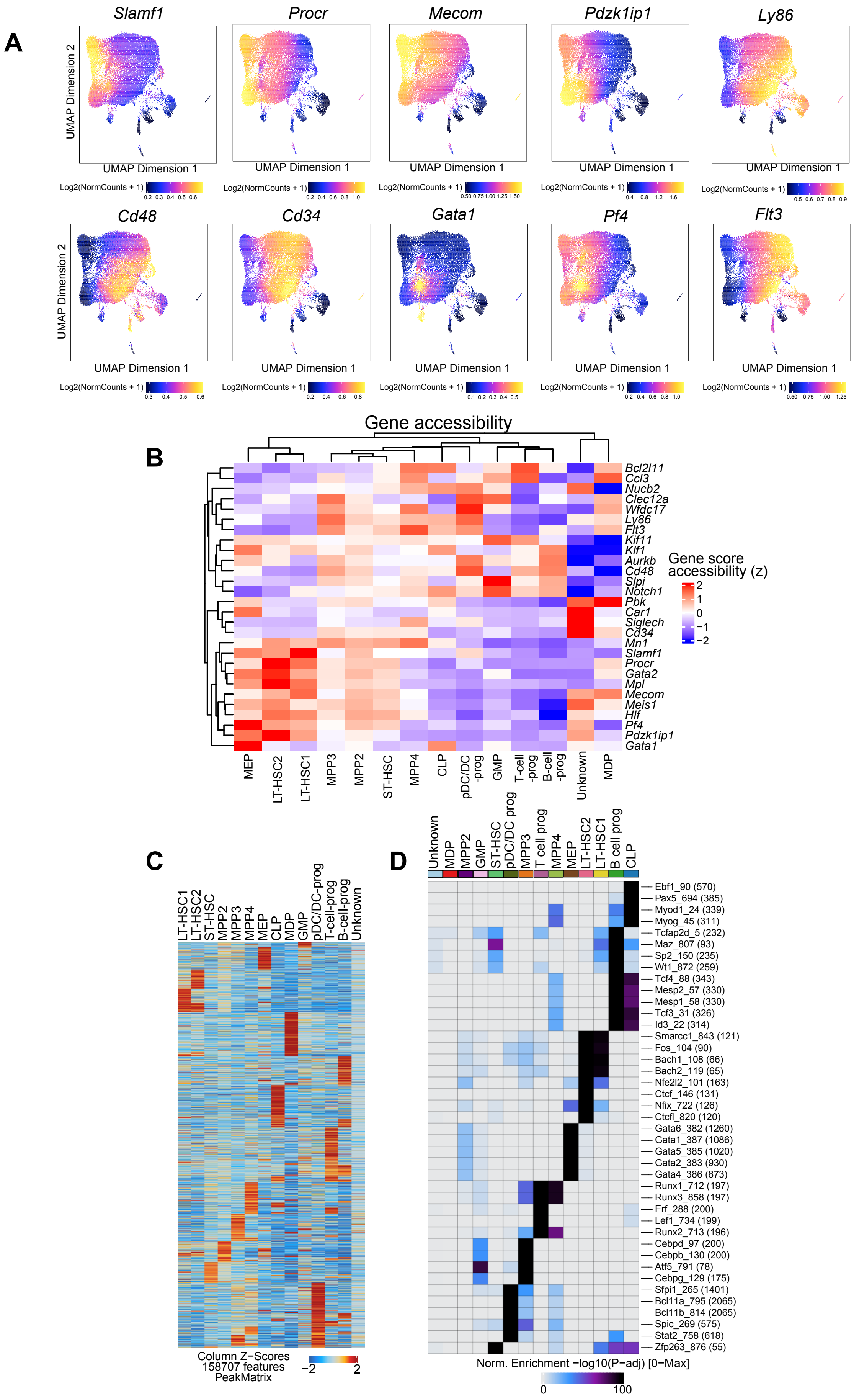

**Figure S6. scATAC-seq cluster identification from LSK cells.**

(**A**) scATAC-seq UMAP-projected gene score accessibility of HSPC cluster marker genes.

(**B**) Heatmap showing quantification of gene score accessibility of HSPC cluster marker genes. Clustering was done using unsupervised clustering for gene accessibility.

(**C**) Heatmap of Z-scores-normalized accessibility of 158,707 differentially accessible peaks in different cell clusters.

(**D**) A heatmap showing -log10 (*P*-adj) value of TF binding motif enrichment in clusters calculated with CIS-BP motif databases, and top 5 motifs with FDR ≤ 0.05 and log2FC ≥ 0.58 are shown

#### Supplemental Figure 7

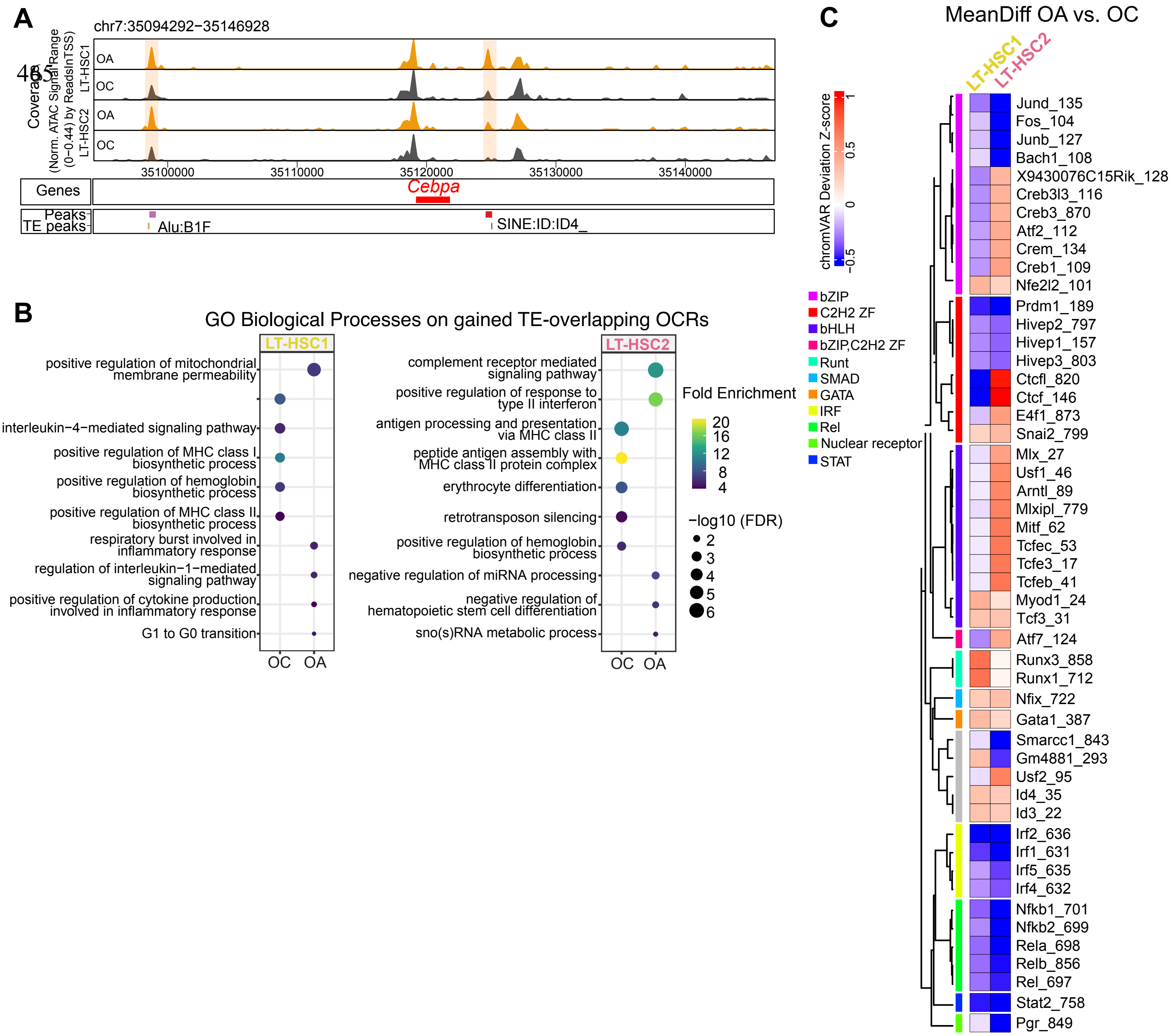

**Figure S7. Chronic alcohol consumption increases chromatin accessibility of TEs.**

**(A)** Browser tracks showing alcohol-induced DA TE peaks correspond to the TEs located approximately 50 kb upstream and downstream of the *Cebpa* in LT-HSC1 and LT-HSC2.

(**B**) Dot plot showing selected GO biological processes of control and alcohol TE-overlapping OCRs. Dot size represents -log10 FDR, color gradation represents Fold Enrichment. The whole list is in **Table S6.**

**(C)** A heatmap showing the mean difference in chromVAR TF deviation Z-scores in LT-HSC clusters OA vs OC. Motifs with FDR $\leq 0.05$ and MeanDiff $\geq 0.58$ calculated using the Wilcoxon test are shown. Columns, but not cell type rows, are hierarchically clustered.

### Supplemental Figure 8

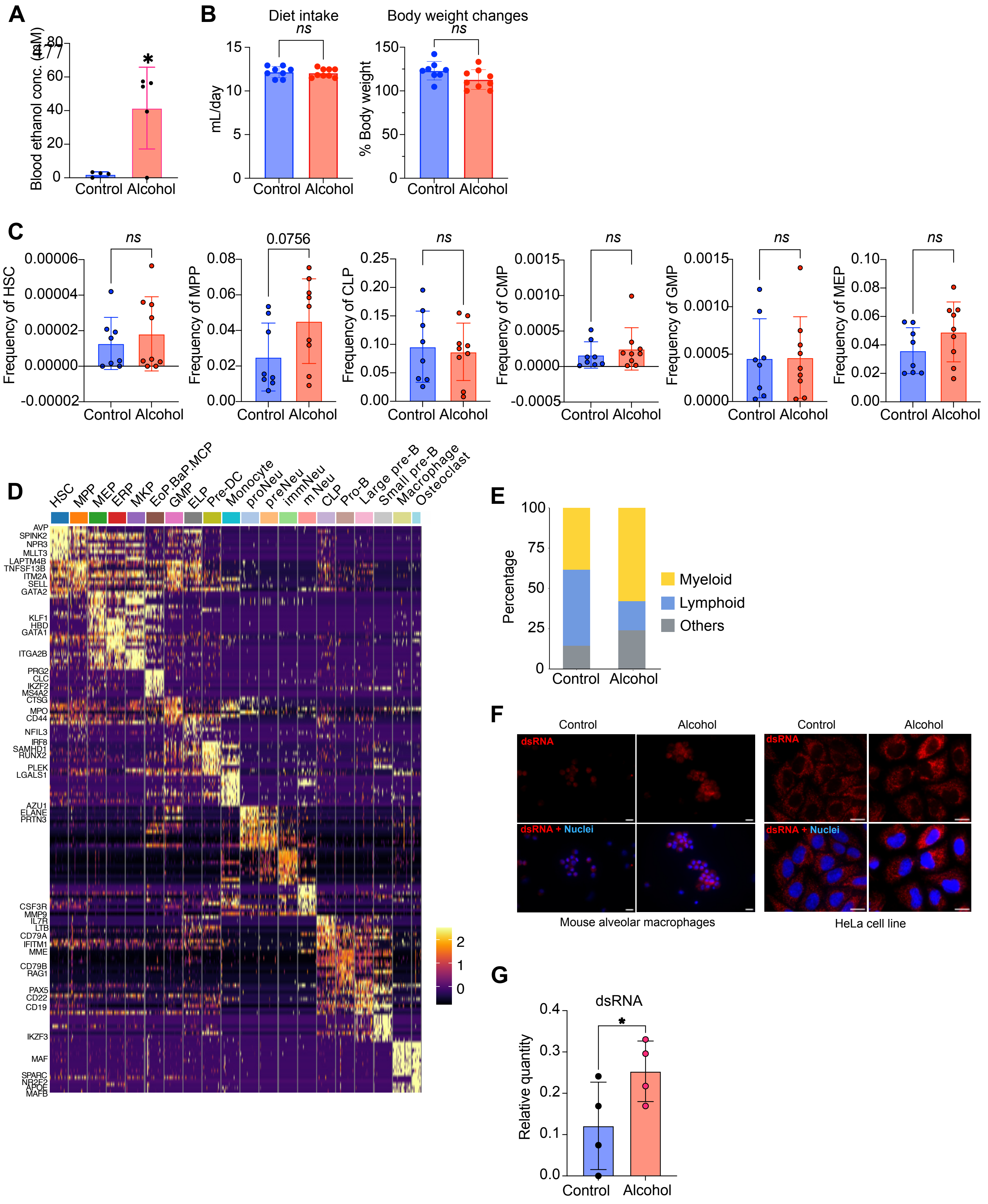

**Figure S8. Chronic alcohol consumption promotes myeloid bias in human CD34+ cells and induces the dsRNA-sensing pathway.**

**(A)** Blood ethanol concentration in mM as measured at the end of 8 weeks of alcohol feeding.

**(B)** Average daily dietary intake and body weight changes at the end of 8 weeks of alcohol feeding.

**(C)** Frequency of several HSPC populations at the end of 8 weeks of alcohol feeding. Data were pooled from two independent experiments. Student's t-test; ns, non-significant. Dots represent individual mice.

**(D)** A heatmap showing the top 20 cluster marker genes calculated using the FindMarkers function and the Wilcoxon rank-sum statistical test with threshold *padj* < 0.01 and absolute log2FC > 1.6.

**(E)** Proportion of myeloid and lymphoid cells in scRNA-seq clusters of human HSPCs from xenotransplant experiments.

**(F)** dsRNA immunofluorescence staining in alveolar macrophages after 10 days of alcohol feeding, and in the HeLa cell line after 4 days of *in vitro* alcohol treatment. Scale bar represents 20 μm**.**

**(G)** Relative concentration of J2 antibody-immunoprecipitated dsRNAs from cultured MOLM13 leukemia cells treated with 100 mM alcohol for 4 days.
